# Subcellular spatial transcriptomics identifies three mechanistically different classes of localizing RNAs

**DOI:** 10.1101/2021.12.05.471303

**Authors:** Lucia Cassella, Anne Ephrussi

## Abstract

Intracellular RNA localization is a widespread and dynamic phenomenon that compartmentalizes gene expression and contributes to the functional polarization of cells. Thus far, mechanisms of RNA localization identified in *Drosophila* have been based on a few RNAs in different tissues, and a comprehensive mechanistic analysis of RNA localization in a single tissue is lacking. Here, by subcellular spatial transcriptomics we identify RNAs localized in the apical and basal domains of the columnar follicular epithelium (FE) and we analyze the mechanisms mediating their localization. Whereas the dynein/BicD/Egl machinery controls apical RNA localization, basally-targeted RNAs require kinesin-1 to overcome a “default” dynein-mediated transport. Moreover, a non-canonical, translation- and dynein-dependent mechanism mediates apical localization of a subgroup of dynein-activating adaptor RNAs (*BicD*, *Bsg25D*, *hook*). Altogether, our study identifies at least three mechanisms underlying RNA localization in the FE, and suggests a possible link between RNA localization and dynein/dynactin/adaptor complex formation *in vivo*.

## INTRODUCTION

RNA localization allows the precise compartmentalization of gene expression in space and time, and is a widespread phenomenon in many different cell types and organisms (Shepard et al., 2003; Blower et al., 2007; Lécuyer et al., 2007; Mili et al., 2008; Jambor et al., 2015; Wilk et al., 2016; Moor et al., 2017). Three main mechanisms have been described to account for RNA localization: (1) active transport on cytoskeletal tracks, (2) localized protection from degradation, or (3) facilitated diffusion and entrapment (Medioni et al., 2012). Recently, several novel mechanisms have been reported to mediate RNA localization, such as hitch-hiking on other RNAs or organelles and co-translational RNA transport (Corradi et al., 2020; Cioni et al., 2019; Liao et al., 2019; Baumann et al., 2014; Harbauer et al., 2021; Cohen et al., 2021; Sepulveda et al, 2018). Active transport is the best characterized mode of RNA localization and consists in the transport of ribonucleoprotein particles by motor proteins on cytoskeletal tracks. Localizing RNAs are typically transported in a translationally silent state and encode cis-acting localization elements (LEs) that are recognized and bound by trans-acting RNA-binding proteins (RBPs) mediating motor recruitment (Xing & Bassell, 2013).

Kinesin motor proteins mostly mediate microtubule (MT) plus end-directed transport. Kinesin-1 (Khc) has been shown to mediate *oskar* (*osk*) RNA localization to the posterior pole of the *Drosophila* oocyte (Brendza et al., 2000; Zimyanin et al., 2008). Whereas Tropomyosin-1 isoform I/C (atypical Tm1, *a*Tm1) regulates *osk* posterior localization by directly stabilizing Khc interaction with the RNA (Dimitrova-Paternoga et al., 2021; Gáspár et al., 2016; Erdélyi et al., 1995), the Exon Junction Complex (EJC) deposited upon splicing is thought to activate kinesin-1 transport of the RNA (Gáspár et al., 2016). Little is known about MT plus end-directed RNA transport in other tissues. Interestingly, *a*Tm1 is also important for *coracle* RNA localization at *Drosophila* neuromuscular junctions (Gardiol & St Johnston, 2014) and the EJC has been shown to mediate *NIN* RNA localization in human RPE1 cells (Kwon et al., 2021).

Cytoplasmic dynein and its accessory complex dynactin direct trafficking of cargoes towards MT minus ends. In *Drosophila*, dynein-mediated RNA transport is accomplished by the dynein-activating adaptor Bicaudal-D (BicD) and the RNA binding protein Egalitarian (Egl) (Mach & Lehmann, 1997; Navarro et al., 2004; Dienstbier et al., 2009). The dynein/BicD/Egl complex is thought to mediate nurse cell-to-oocyte transport of maternal RNAs, and was shown to direct apical RNA localization in the early embryo, neuroblasts, and polar cells (Clark et al., 2007; Wilkie & Davis, 2001; Bullock & Ish-Horowicz, 2001; Hughes et al., 2004; Van De Bor et al., 2011). The dynein/dynactin/BicD (DDB) motor complex is highly conserved and participates in the transport of different cargoes, with BicD (and its mammalian ortholog BICD2) linking the dynein motor to specific cargoes. While proteins binding to the BicD C-terminal domain (CTD), such as Egl or Rab6, impart cargo specificity (Matanis et al., 2002; Hoogenraad et al., 2003; Dienstbier et al., 2009; Coutelis & Ephrussi 2007; Januschke et al., 2007), the BicD N-terminal domain (corresponding to coiled-coil 1/2, CC1/2) binds to dynein/dynactin (Hoogenraad et al., 2001, 2003) and activates dynein processivity (McKenney et al., 2014; Schlager et al., 2014; Dienstbier et al., 2009; Sladewski et al., 2018).

Although much of what is known about RNA localization comes from studies of maternally inherited RNAs in the *Drosophila* germline, several examples of localizing RNAs have been also reported in the follicular epithelium (FE) that envelops the germline cyst (Jambor et al., 2015; Li et al., 2008; Horne-Badovinac & Bilder, 2008; Vazquez-Pianzola et al., 2017; Schotman et al., 2008; Serano & Rubin, 2003). The FE is composed of highly polarized secretory follicle cells (FCs) belonging to the somatic lineage, with minus ends of non-centrosomal microtubules (ncMTs) anchored at the apical cell cortex facing the oocyte (Clark et al., 1997; Khanal et al., 2016). The FE is an easily manipulatable and powerful genetic system that, through the generation of mosaics, allows the dissection of the effect of mutations without disrupting developmental processes. Several lines of evidence indicate that the dynein/BicD/Egl RNA transport complex active in nurse cell-to-oocyte transport is also responsible for the apical localization of a handful of RNAs in the FE (Li et al., 2008; Bhagavatula & Knust, 2021; Karlin-McGinness et al., 1996; Jambor et al., 2014; Vazquez-Pianzola et al., 2017; Van De Bor et al., 2011). However, a comprehensive overview of RNA localization in the FE and its underlying mechanisms are lacking.

Here, we apply subcellular spatial transcriptomics to first identify the landscape of apically- and basally-localizing RNAs in the columnar FE. By screening a subset of apical and basal RNAs identified in this way, we find that the dynein/BicD/Egl machinery acts by “default” in directing apical RNA localization, and that an additional kinesin-1-dependent layer of regulation must be applied to direct basal RNA localization. Moreover, we identify a third, translation- and dynein-dependent mechanism that underlies the apical localization of transcripts encoding dynein-activating adaptors, providing a possible link between RNA localization and dynein/dynactin/adaptor complex formation *in vivo*.

## RESULTS

### Identification of apical and basal RNAs in columnar follicle cells/bold>

To identify RNAs that localize apically or basally in *Drosophila* FE transcriptome-wide, we applied laser-capture microdissection (LCM) to isolate fragments of tissue that consisted in either the apical half (“apical domain”) or basal half (“basal domain”) of adjacent columnar follicle cells (**Figure 1A and Movie S1**). Differential gene expression analysis of apical vs. basal LCM-derived RNA-seq samples yielded 306 RNAs enriched in the apical samples and 249 RNAs enriched in the basal samples (false discovery rate [FDR] < 0.1) (**Figure 1B**). Since LCM is highly susceptible to tissue contamination, we first aimed at identifying those RNAs whose significant enrichment was a result of contamination by other cell types, such as the oocyte on the apical side or the circular muscles on the basal side (**Figure S1A**). To do so, we analyzed those RNAs characterized by high absolute log2-transformed fold change (|log2FC|) values of apical over basal abundance that might result from contamination of neighboring tissues expressing a different set of hallmark genes. By setting an arbitrary threshold of |log2FC| > 3 as indicative of contaminant RNA identity, we found 33 putative basal contaminants of muscle origin (log2FC < −3) and 2 putative apical contaminants of oocyte origin (log2FC > 3) (**Figure 1B** and **Figure S1B**). 2/3 (n=22) of basal genes with log2FC < −3 were annotated as being expressed or having a function in muscle tissues (FlyBase) and their mapped reads were often absent or in very low number in the apical fragments (**Figure S1C,D**). Moreover, we validated through single molecule Fluorescence *In Situ* Hybridization (smFISH) 3 putative basal contaminants (*Mhc*, *Act57B*, *wupA*) as being enriched in circular muscles with little or no expression in the FE (**Figure S1E**). This analysis resulted in 304 *bona fide* apical RNAs and 216 *bona fide* basal RNAs localizing in the columnar FE (**Figure 1B, Table S1**). Finally, 16 RNAs were randomly chosen from the computationally established list of significantly enriched *bona fide* apical or basal RNAs and were validated as true localizing RNAs through smFISH (**Figure 1C**).

**Figure 1.**
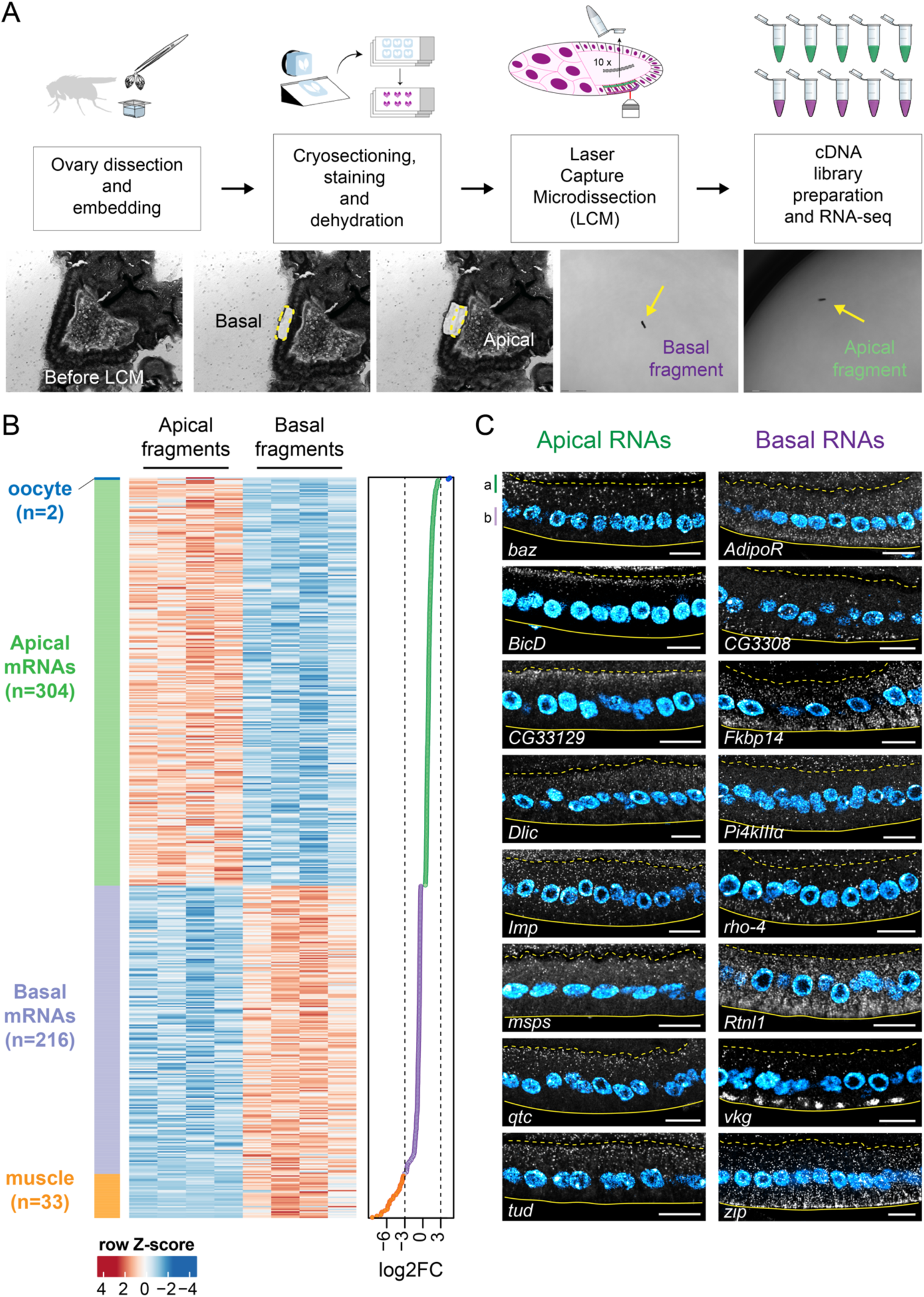
**Identification of apical and basal RNAs in *Drosophila* follicular epithelium by subcellular spatial transcriptomics.** A) Schematic representation of the sample preparation procedure. Lower panels: representative images of an egg chamber before and after apical and basal fragment microdissection, including visualization of microdissected fragments in the cap of collection tubes. B) Heatmap representing RNA-seq signal (*z*-score of normalized read counts) for significantly enriched RNAs in microdissected apical and basal fragments (FDR < 0.1). Each row represents a significantly enriched RNA in either apical samples (n=4) or basal samples (n=4). The log2FC value of each RNA showed in the heatmap is indicated in the graph on the right. Dashed lines indicate threshold log2FC values (log2FC < −3 and log2FC > 3) arbitrarily set to identify oocyte contaminants (log2FC > 3, n=2, blue), *bona fide* apical RNAs (0 < log2FC ≤ 3, n=304, green), *bona fide* basal RNAs (−3 ≤ log2FC < 0, n=216, purple), and muscle contaminants (log2FC ≤ −3, n=33, orange). C) smFISH validation of 16 *bona fide* apical (left panels) and basal (right panels) RNAs. A dashed line and a continuous line in each panel delimit the FC-oocyte and FC-basal lamina borders respectively. a = apical domain; b = basal domain. Nuclei (cyan) are stained with DAPI. Scale bars 10 μm. See also Figure S1, Table S1 and Video S1.

### Basal RNA localization depends on kinesin-1, *a*Tm1, and the EJC

Basal RNA localization is a largely uncharacterized phenomenon. Previous reports have identified a limited number of basally-localizing RNAs in the FE (Jambor et al., 2015; Schotman et al., 2008; Serano & Rubin, 2003), with little mechanistic insight. For this reason, we sought to elucidate the mechanisms behind basal RNA localization. Early reporter-based studies on the polarity of *Drosophila* tissues have shown that the basal domain of the FE is functionally equivalent to the posterior pole of the oocyte, as both compartments accumulate the MT plus end marker Kin:βgal (Clark et al., 1997).

Therefore, we hypothesized that the regulators of *oskar* posterior RNA transport might also be responsible for basal RNA localization in the FE. To test this hypothesis, we disrupted known components of the *osk* RNP transport machinery, such as kinesin-1 (Khc), atypical Tropomyosin-1 (*a*Tm1) and the Exon Junction Complex (EJC) (**Figure S2A)** in the FE and analyzed the localization pattern of 4 validated basal RNAs (*Fkbp14*, *CG3308*, *Rtnl1*, *zip*) (**Figure 2**). In all cells lacking either Khc (*Khc* RNAi cells) (**Figure 2A**), *a*Tm1 (*Tm1^NULL^*, Erdélyi et al., 1995) (**Figure 2B**), or the EJC (*ΔC-Pym* cells, Ghosh et al., 2014) (**Figure 2C**), basal RNA localization was severely disrupted, with all basal RNAs analyzed becoming apically localized. To check whether the changes observed in RNA localization were specific of basal RNAs, we analyzed the localization pattern of 4 apical RNAs validated previously (*crb*, *msps*, *qtc, CG33129*) in the same mutants. In contrast to basal RNAs, none of the apical RNAs analyzed were affected by disruption of kinesin-1-mediated RNA transport (**Figure S2B-D**), indicating that regulators of RNA transport towards MT plus ends specifically control basal RNA localization. To have a quantitative overview of changes in RNA localization, we considered the ratio between the apical and the basal smFISH signal intensity in either wild-type (wt) or knock-down (KD) cells, and called this parameter Degree of Apicality (DoA), as values > 1 indicate an apical localization bias. Then, we tested whether the DoA values of each RNA analyzed significantly differ in KD vs. wt cells by calculating the ratio between the DoA(KD) and the DoA(wt) for each RNA in each of the 3 conditions (see Materials and Methods and Figure 2 for statistical testing). With this analysis, we confirmed that (1) all basal RNAs were affected by lack of Khc, *a*Tm1, or EJC and (2) none of the apical RNAs significantly changed localization pattern upon knock-down of kinesin-1 regulators (**Figure 2D**), showing that kinesin-1, *a*Tm1, and the EJC are specifically responsible for basal RNA localization in the FE.

**Figure 2.**
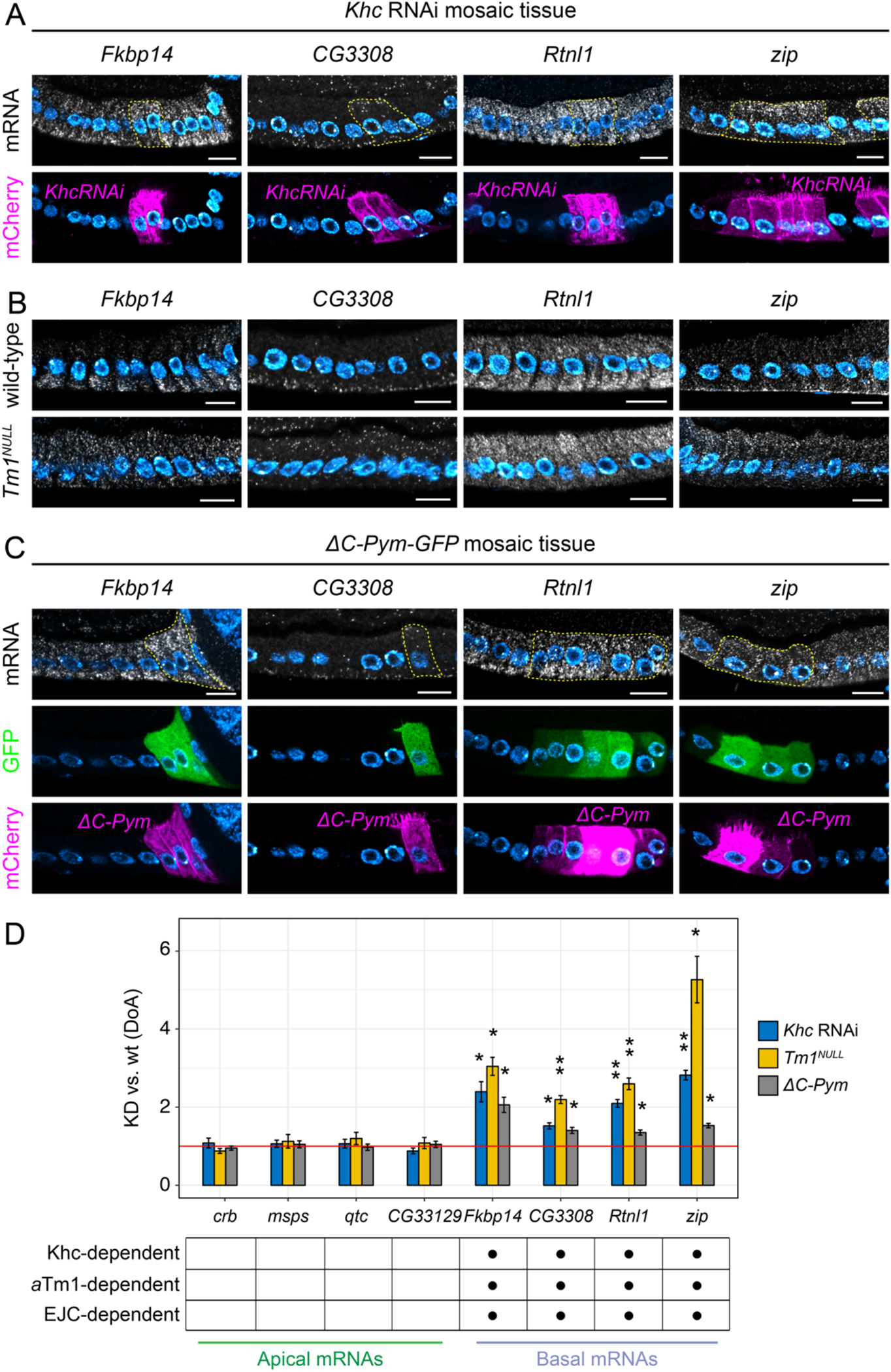
**Basal RNA localization depends on kinesin-1, *a*Tm1, and the EJC.** In A) and C), mutant cells (marked with CD8-mCherry, lower panels) were generated by the UAS/Gal4 FLP-out system by inducing *Khc* RNAi (A) or by expressing the EJC-disrupting protein *ΔC-Pym* (C), to disrupt each component without significantly affecting tissue architecture. Neighboring wild-type cells are unmarked. A dashed line highlights mutant cells in smFISH images (upper panels). In B) the expression of the *a*Tm1 isoform was specifically knocked down by generating *Tm1^eg9^/Tm1^eg1^* (*Tm1^NULL^*) egg chambers. A) Localization of basal RNAs by smFISH in *Khc* RNAi mosaic tissue. B) Localization of basal RNAs by smFISH in wild-type and *Tm1^NULL^* egg chambers. C) Localization of basal RNAs by smFISH in *ΔC-Pym-GFP* mosaic tissue. D) Quantification of changes in the A-B distribution of apical and basal RNAs in conditions of downregulated kinesin-1 transport. Analyzed RNAs are indicated on the x-axis. The y-axis shows the average values (± s.e.m) of the ratio between the Degree of Apicality (DoA) measured in knock-down (KD) cells and the DoA measured in wild-type (wt) cells for each RNA analyzed, in each of the three conditions. The mean KD/wt(DoA) value for each RNA in each condition was tested against a null hypothesis *H_0_* of KD/wt(DoA)=1 (red horizontal line), corresponding to no change between mutant and wild-type cells (one-sample t-test). Asterisks indicate mean values that significantly differ from the reference value of mu=1 (*=p<0.05; **=p<0.01; ***=p<0.001). Nuclei (cyan) are stained with DAPI. Scale bars 10 μm. See also Figure S2 and Figure S3.

### Mislocalization of the basal RNA *zip* depends on Egalitarian

Interestingly, upon disruption of MT plus end-directed RNA transport all analyzed basal RNAs were mislocalized to the apical domain. Several studies reported that apical RNA localization depends on the BicD/Egl machinery, a dynein-dependent complex that localizes RNAs apically in the blastoderm embryo and is thought to be responsible for nurse cell-to-oocyte transport of maternal RNAs. Therefore, the apical mislocalization of basal RNAs observed upon knock-down of kinesin-1 regulators might be due to apical RNA transport by the dynein/BicD/Egl machinery. To test this, we generated FC clones lacking either Egl (*egl* RNAi) or Khc (*Khc* RNAi), or both Egl and Khc [(*egl*+*Khc*) RNAi] and evaluated changes in the RNA localization of *zip*, one of the most striking examples of the apical mislocalization phenomenon (see Figure 2A-C). Whereas *zip*RNA was unaffected upon *egl* RNAi and strongly apically mislocalized in *Khc* RNAi conditions as also highlighted by our previous experiments, (*egl*+*Khc*) RNAi caused *zip* to assume a ubiquitous localization that would be consistent with a failure of both kinesin-1-and dynein-mediated transport (**Figure S3A**). *zip*DoA measurements in wt and *RNAi* cells in each of the three conditions provided a quantitative evaluation of the changes observed in smFISH experiments (**Figure S3B**), with a significant decrease in KD/wt DoA in double (*egl+Khc*) RNAi cells (KD/wt DoA = 1.61) compared to *Khc* RNAi cells (KD/wt DoA = 2.49) (**Figure S3C**).

Therefore, despite being dispensable in basal RNA localization under normal conditions, the dynein/BicD/Egl complex is responsible for the apical mislocalization of a basal RNA (and possibly more) when kinesin-1 activity is lacking.

### Two different dynein-dependent mechanisms control apical RNA localization

As mentioned previously, several reports have identified the dynein/BicD/Egl machinery as responsible for the apical localization of a subset of RNAs in the FE, such as *crumbs* (*crb*) (Li et al., 2008; Bhagavatula & Knust, 2021). To test in an unbiased way the degree of involvement of the dynein/BicD/Egl machinery in the localization of apical RNAs in the FE, we generated FC mutant clones in which either cytoplasmic dynein (*Dhc64C*, hereafter called *Dhc*) or Egalitarian (*egl*) were knocked-down by RNAi (**Figure S4A**). We then analyzed the localization pattern of 5 validated apical RNAs (*crb*, *msps*, *qtc, CG33129*, *BicD,* with *crb* RNA as a positive control) by smFISH, along with the quantification of RNA localization by measuring the KD/wt DoA. The localization of all apical RNAs analyzed was completely abolished when *Dhc* was knocked down by RNAi, with the RNAs becoming ubiquitously distributed (**Figure 3A,C**). *egl* RNAi caused all apical RNAs to lose their apical localization, with the surprising exception of *BicD* (**Figure 3B,C**; see below). In contrast, basal RNAs largely maintained their basal localization pattern upon either *Dhc* RNAi or *egl* RNAi treatment (**Figure S4B,C**and **Figure 3C**). Basal RNA localization was only mildly affected in a subset of Dhc RNAi cells, likely as a consequence of the emergence of polarity defects in cells lacking Dhc (Horne-Badovinac & Bilder, 2008; Ronchi et al, 2021) (see Figure 3A and Figure S3B). The maintenance of BicD RNA localization in egl RNAi cells was not due to a low efficiency of the RNAi, since both egl RNA and Egl protein were significantly reduced in egl KD cells (**Figure S4D,E**). Moreover, in egg chambers entirely lacking Egl throughout the FE (eglNULLFC, see Materials and Methods), BicD RNA was still apically localized, whereas localization of CG33129 RNA, previously found to be Egl-dependent (see Figure 3B), was disrupted (**Figure S4F**). Altogether, these results show that the dynein/BicD/Egl complex is largely responsible for apical RNA localization, but a different dynein-dependent mechanism underlies the apical localization of BicD RNA. Considering that the Egl-independent targeting of BicD RNA represents a novel mechanism of apical RNA localization, we sought to gain more insight into the mechanisms regulating its RNA transport.

**Figure 3.**
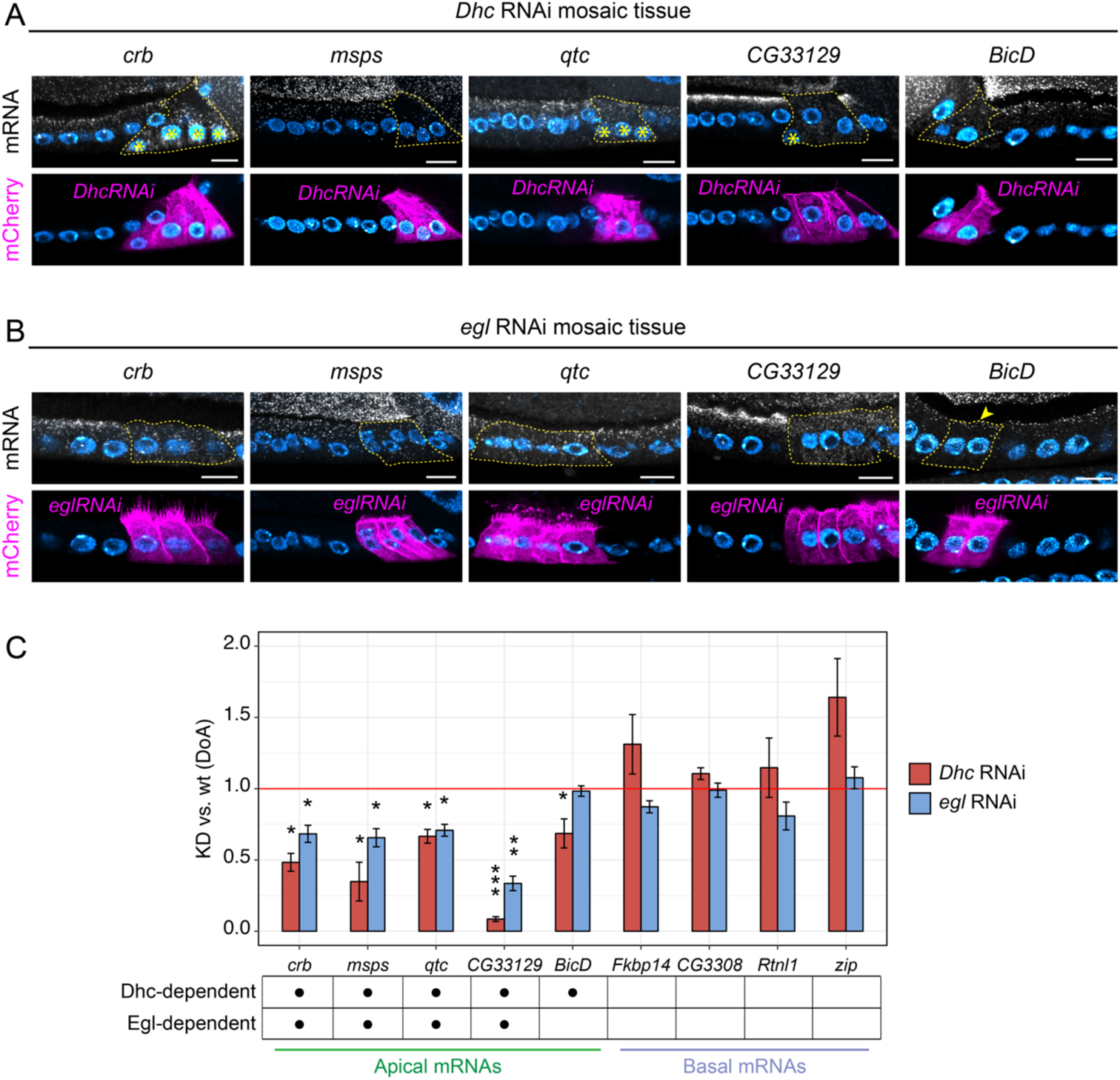
**Two different dynein-dependent mechanisms control apical RNA localization.** A-B) Localization of apical RNAs by smFISH in *Dhc* RNAi (A) and *egl* RNAi mosaic tissue (B). Mutant cells are marked by the expression of CD8-mCherry (lower panels) and highlighted with a dashed line in smFISH images (upper panels). Neighboring wild-type cells are unmarked. Asterisks (*) indicate basal mispositioning of nuclei due to *Dhc* RNAi, an indication of mild cell polarity defects. The arrowhead in B) indicates the persistence of apical *BicD* RNA in *egl* RNAi cells. C) Quantification of changes in the A-B distribution of apical and basal RNAs in conditions of downregulated dynein/BicD/Egl transport (*Dhc* RNAi or *egl*RNAi). Analyzed RNAs are indicated on the x-axis. The y-axis shows the average values (± s.e.m) of the KD/wt ratio (DoA) for each RNA analyzed, in each of the two conditions. The mean KD/wt(DoA) value for each RNA in each condition was tested against a value of KD/wt(DoA)=1 (red horizontal line), corresponding to no change between mutant and wild-type cells (one-sample t-test). Asterisks indicate mean values that significantly differ from the reference value of mu=1 (*=p<0.05; **=p<0.01; ***=p<0.001). Nuclei (cyan) are stained with DAPI. Scale bars 10 μm. See also Figure S4.

### *BicD* RNA localization requires an intact translation machinery

Localization of *BICD2/BicD*RNA at centrosomes in cultured cells is translation-dependent (Safieddine et al., 2021). To test whether *BicD* RNA localization in the FE involves the same mechanism, we treated egg chambers *ex vivo*with the translation inhibitors puromycin (Puro) and cycloheximide (CHX) and analyzed the distribution of *BicD*RNA under these two conditions compared to control ovaries incubated in Schneider’s medium only (**Figure 4A**). To assess tissue integrity, in parallel we visualized *osk* RNA, whose localization during the middle stages of oogenesis should not be affected by translation inhibitors. Whereas the localization pattern of *BicD* RNA in CHX-treated egg chambers was similar to controls (**Figure 4B,D**), Puro treatment clearly impaired *BicD* RNA localization in the FE (**Figure 4C**). The distribution of *BicD* signal intensity along the A-B axis of mid-stage follicle cells shows that *BicD* enrichment at the apical cortex of the FE was severely reduced upon Puro treatment (**Figure 4E**). As a proxy for the degree of signal mislocalization, we calculated the value corresponding to 50% of the cumulative area under the curve (a.u.c.) in Puro- or CHX-treated egg chambers and compared it with untreated controls. The results of this analysis show that the *BicD* RNA signal shifted significantly towards the basal domain in Puro-treated egg chambers, whereas CHX had no effect on *BicD* RNA localization (**Figure 4F**). The fact that freezing elongating ribosomes (CHX condition) does not affect *BicD* RNA localization, whereas blocking translation by releasing the nascent peptide (Puro condition) does, indicates that an intact translation machinery and the presence of a nascent peptide may be required for *BicD* RNA localization in FCs.

**Figure 4.**
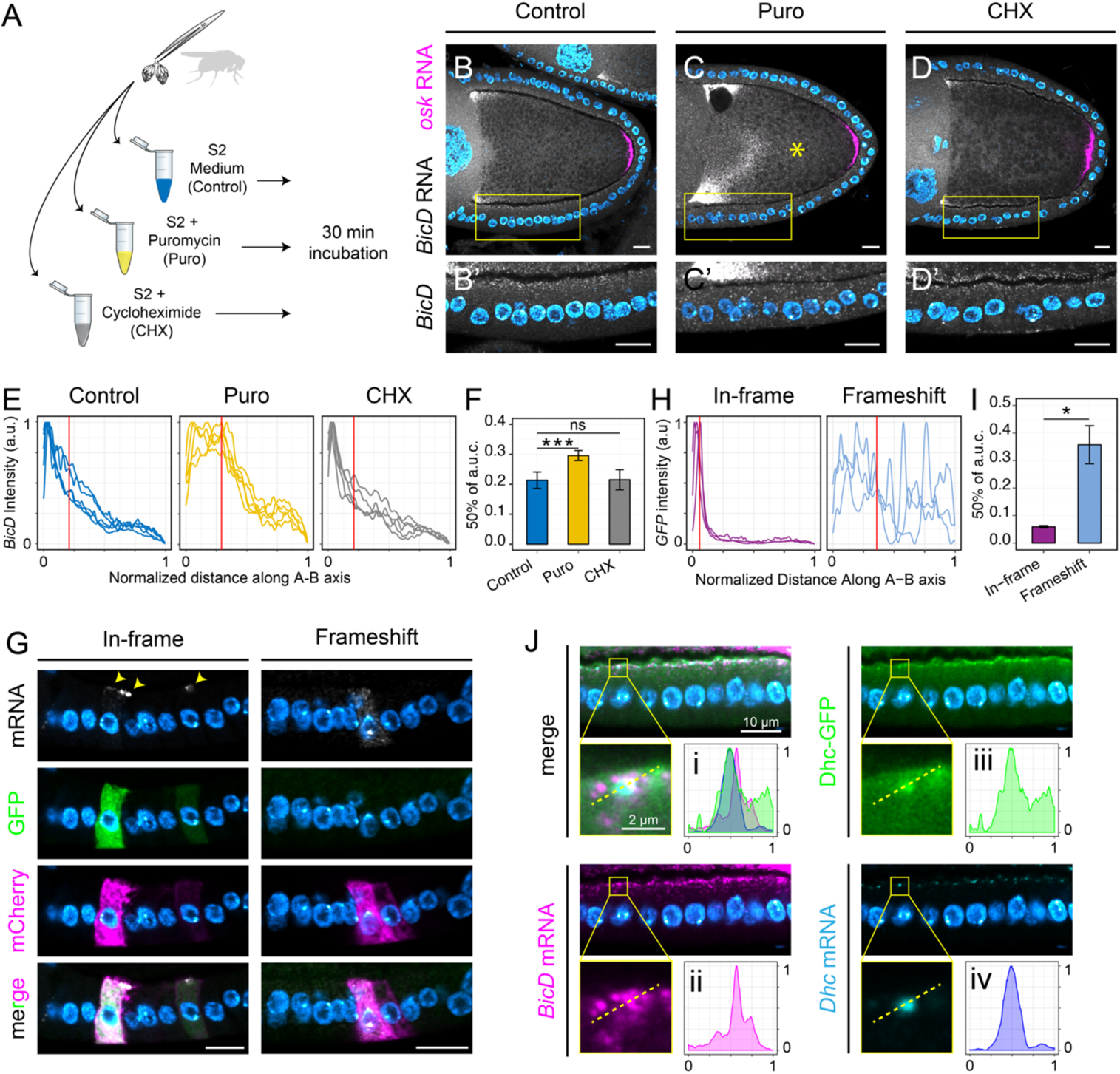
***BicD* RNA is co-translationally localized at cortical Dhc foci.** A) Schematic representation of the *ex vivo* treatment of wild-type ovaries with pharmacological inhibitors of translation. B-D) Dual-color *BicD* (grayscale) and *osk* (magenta) smFISH experiments on Control (B-B’), Puro-(C-C’) or CHX-treated (D-D’) ovaries. Insets show a magnification of the follicular epithelium (bottom panels). Note the mislocalization of *BicD* RNA towards the center of the oocyte (*) in C). E) Quantification of *BicD* RNA distribution along a linear ROI spanning the apical-basal axis of follicle cells measured as smFISH fluorescence intensity in control (blue), Puro (yellow) and CHX (grey) conditions. A red vertical line represents the mean x value corresponding to the 50% of the cumulative area under the curve (a.u.c.), a proxy for *BicD* degree of mislocalization. F) Statistical analysis of *BicD* degree of mislocalization (mean a.u.c. ± s.e.m) in each condition compared to control. Control-Puro: p=0.000879 (***); Control-CHX: p=0.9374 (ns). G) Expression of different *BicD-GFP* constructs (“*In-frame*”: *^0^BicD-GFP*; “*Frameshift*”: *^(−1)^BicD-GFP* or *^(+1)^BicD-GFP*) in FC clones and analysis of transgenic RNA distribution by smFISH with antisense GFP probes. Follicle cell clones expressing each *BicD-GFP* construct are marked by CD8-mCherry (magenta). H) Quantification of *In-frame* (purple) or *Frameshift* (light blue) *BicD-GFP* RNA distribution. I) Statistical analysis of *BicD-GFP* degree of mislocalization (mean a.u.c. ± s.e.m) in *Frameshift* compared to *In-frame* construct. p= 0.01719 (*). J) Localization of *BicD* RNA (magenta), *Dhc* RNA (cyan), and endogenously tagged Dhc-GFP (green) in stage 10 follicular epithelium. Insets show a magnification of a single Dhc-GFP/*Dhc* RNA focus. A dashed line indicates the cross-section along which each signal was measured (panels i-iv). Signal intensities (y-axis) and line length (x-axis) were normalized in the 0-1 range. Nuclei (cyan) are stained with DAPI. Scale bars 10 μm unless otherwise specified. See also Figure S5 and Figure S6.

### *BicD* RNA is co-translationally localized

To understand whether the localization of *BicD* depends on translation of its own RNA (in *cis*) or of other factors (in *trans*), we designed a series of transgenic constructs consisting of a BicD-GFP cassette inserted downstream of an 18-bp linker in which we could introduce the desired frameshift mutations without disrupting any unknown RNA localization element in the BicD CDS (**Figure S5A**). Each of these transgenes was expressed in FC clones in a *BicD* wild-type background and the transgenic *BicD-GFP* RNA was specifically detected by smFISH using antisense GFP probes. *GFP* RNA carrying the same 3’ untranslated region (UTR) as BicD-GFP constructs failed to localize when expressed in the germline or in the FE (**Figure S5B**), showing that this sequence alone is not sufficient to drive RNA localization. In contrast, the in-frame *^0^BicD-GFP* RNA showed a strong apical localization in FCs (**Figure 4G-I**), similarly to the endogenous *BicD* RNA (see Figure S6A). Moreover, the expression of full-length BicD-GFP was validated by the presence of GFP fluorescence in CD8-mCherry^+^ cells expressing the transgene (**Figure 4G**). Disruption of the BicD-GFP reading frame by either +1 or −1 frameshift, verified by the absence of GFP signal in CD8-mCherry^+^ cells, was sufficient to impair apical RNA localization (**Figure 4G-I**). Consistent with the puromycin-induced impairment of RNA localization in the FE, these results show that *BicD*RNA is co-translationally localized at the apical cortex.

### *BicD* and *Dhc* RNAs decorate dynein particles at the apical cortex

As in BicD the first peptide emerging from the ribosome is the dynein-binding domain (Hoogenraad et al., 2003), the co-translational localization of *BicD* RNA might depend on association of nascent BicD protein with dynein. To have an indication whether this might be the case, we imaged *BicD* RNA by smFISH in ovaries expressing endogenously tagged Dhc-GFP (Gaspar et al., 2021). Although the Dhc-GFP signal was diffuse in the ovary, distinct Dhc-GFP foci were detected at the apical cortex of columnar FCs (**Figure 4J**) and elsewhere in the germline (see below). These foci also contain *Dhc* RNA, indicating that these might be sites of *Dhc* RNA translation. *BicD*RNA showed a partial co-localization with Dhc-GFP/*Dhc* RNA foci, consistent with the hypothesis of its co-translational association with newly synthesized Dhc protein at the apical cortex.

### The first step of *BicD* RNA localization in the early cyst is translation-independent

In the germline, BicD has an instructive role in oocyte specification (Wharton & Struhl, 1989; Suter & Steward, 1991; Mach & Lehmann, 1997). Importantly, *BicD*RNA localization reflects MT minus end enrichment (Steinhauer & Kalderon, 2006; Clark et al., 1997) in both the germline and FE (**Figure S6A**). We noticed that, as in the FE, *BicD* RNA localization to the posterior of the oocyte (stages 9-10) was impaired in Puro-treated ovaries, whereas CHX had no effect (**Figure 4C,D**). The same effect was visible in younger egg chambers, starting when BicD becomes posteriorly localized in the small oocyte at stages 4-5 (**Figure S6B**). In contrast, neither Puro nor CHX treatment abolished *BicD* nurse cell-to-oocyte transport in early egg chambers, with *BicD* enrichment in the small oocyte being close to wild-type levels (**Figure S6B**). Consistent with this, germline-driven *Frameshift* BicD-GFP RNA underwent nurse cell-to-oocyte transport and displayed a clear oocyte enrichment during early stages, similarly to endogenous *BicD* (**Figure S6C**). However, within the oocyte, *Frameshift* RNA was ubiquitously distributed at these stages, and failed to localize at the posterior cortex of the small oocyte. These results indicate that the process of *BicD* RNA transport into the oocyte does not involve active translation; on the other hand, *BicD* RNA localization within the oocyte is likely governed by the same co-translational mechanism that operates in the FE. In support of this hypothesis, we found that *BicD*RNA decorates Dhc/*Dhc* RNA foci in both the FE and the oocyte, but not in the nurse cells (**Figure S6D**). Taken together, these results indicate that the mid-oogenesis oocyte and the columnar FCs share a similar co-translational mechanism for *BicD* RNA localization. In contrast, *BicD* RNA nurse cell-to-oocyte localization appears to be mediated by a translation-independent mechanism that does not involve the association with Dhc/*Dhc* RNA particles.

### A subset of dynein-activating adaptor RNAs are also co-translationally localized in the FE

BicD belongs to the class of dynein-activating adaptors, linking cargoes to the dynein motor complex (Olenick & Holzbaur, 2019). We found that the RNA encoding all *Drosophila* orthologs of the currently known or putative dynein activating adaptors (hereafter collectively called “adaptor RNAs”), namely *hook* (HOOK2-3), *Bsg25D* (NIN/NINL), *Nuf* (RAB11FIP3), and *Milton* (TRAK1-2), were significantly enriched apically in our list of localizing transcripts (**Table S1**), with the exception of *Spindly* (SPDL1) which was below the detection threshold. By hypothesizing that the same dynein-dependent co-translational process that drives *BicD*RNA localization would also be responsible for the apical localization of adaptor RNAs, we tested whether the localization of adaptor RNAs was affected by either *Dhc* or *egl* RNAi. With the exception of *Nuf* and *Milton*, whose localization was disrupted by either treatment (data not shown), the apical localization of *Bsg25D*and *hook* (**Figure 5A**) was significantly disrupted in *Dhc* RNAi cells (**Figure 5B,D**), but not in *egl* RNAi cells (**Figure 5C,D**).

**Figure 5.**
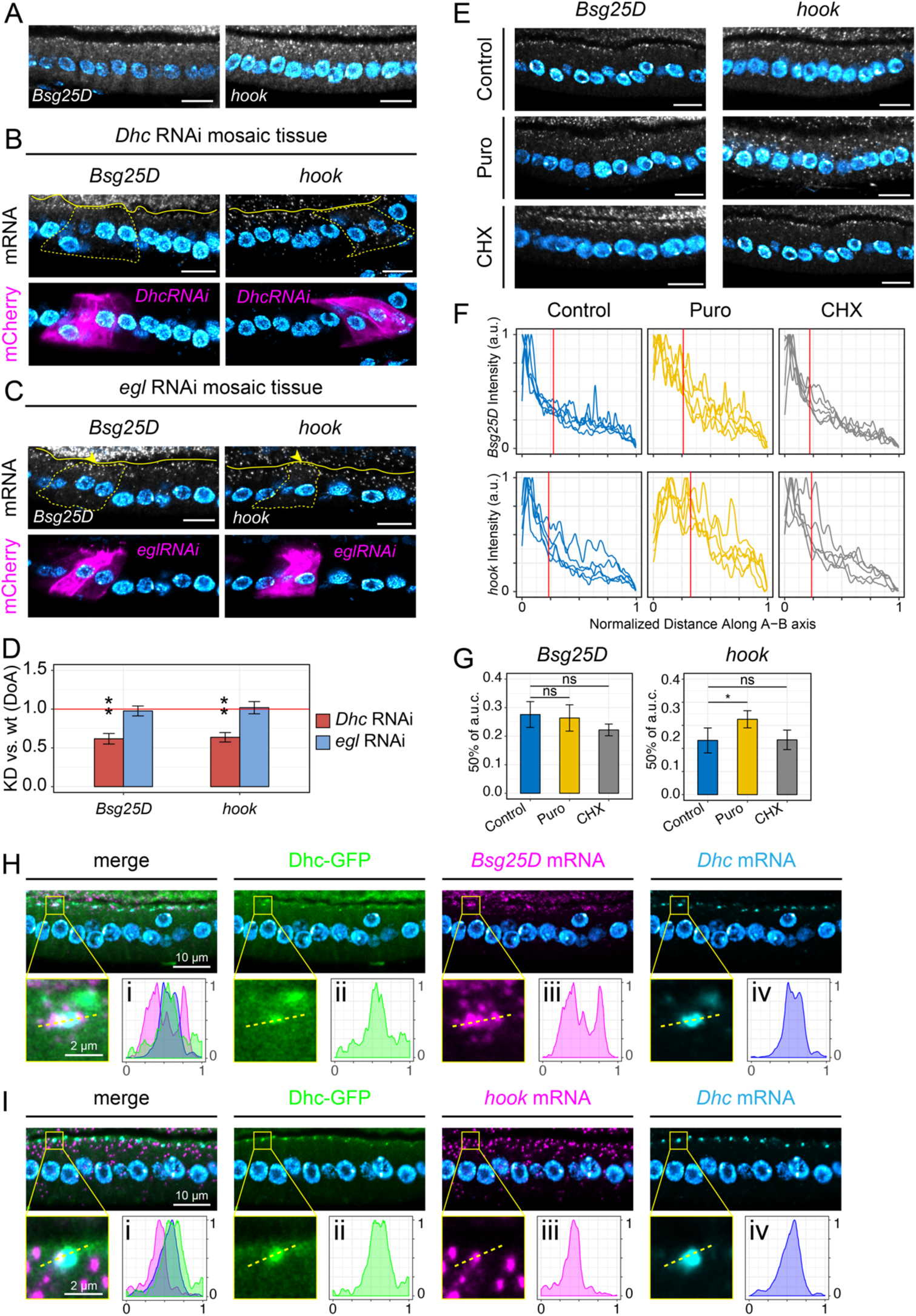
**The apical localization of *hook* and *Bsg25D* RNAs is mechanistically similar to *BicD*.** A) *Bsg25D* and *hook* RNAs visualized by smFISH in wild-type FE. B-C) Localization of *Bsg25D* and *hook* RNAs by smFISH in *Dhc* RNAi (B) and *egl*RNAi mosaic tissue (C). Mutant cells are marked by the expression of CD8-mCherry (lower panels) and highlighted with a dashed line in smFISH images (upper panels). Neighboring wild-type cells are unmarked. Arrowheads in C) indicate the persistence of *Bsg25D* and *hook* RNAs apically in *egl* RNAi cells. A continuous yellow line demarcates the border between the oocyte and the FE to facilitate the image interpretation. D) Quantification of changes in the A-B distribution of *Bsg25D* and *hook* RNAs in conditions of downregulated dynein/BicD/Egl transport. The y-axis shows the average values (± s.e.m) of the KD/wt DoA for each RNA analyzed, in each condition. The mean KD/wt DoA value for each RNA in each condition was tested against a value of KD/wt(DoA)=1 (red horizontal line). Asterisks indicate mean values that significantly differ from the reference value of mu=1 (*=p<0.05; **=p<0.01; ***=p<0.001). E) *Bsg25D*(left panels) and *hook* (right panels) RNA localization in the FE visualized by smFISH in Control, Puro- or CHX-treated ovaries. E) Quantification of *Bsg25D* (upper panels) and *hook* (lower panels) RNA distribution along linear ROIs spanning the apical-basal axis of follicle cells measured as smFISH fluorescence intensity in control (blue), Puro (yellow) and CHX (grey) conditions. A red vertical line represents the mean x value corresponding to 50% of each area under the curve (a.u.c.). G) Statistical analysis of *Bsg25D* and *hook* degree of mislocalization (mean a.u.c. ± s.e.m) in each condition compared to control. Control-Puro(*Bsg25D*): p= 0.6869 (ns); Control-CHX(*Bsg25D*): p=0.05405 (ns), Control-Puro(*hook*): p= 0.01648 (*); Control-CHX(*hook*): p= 0.9412 (ns). H-I) Localization of *Bsg25D* RNA (H) or *hook* RNA (I) (magenta), *Dhc* RNA (cyan), and endogenously tagged Dhc-GFP (green) in stage 10 follicular epithelium. Insets show a magnification of a single Dhc-GFP/*Dhc* RNA focus. A dashed line indicates the cross-section along which each signal has been measured (panels i-iv). Signal intensities (y-axis) and line length (x-axis) were normalized in the 0-1 range. Nuclei (cyan) are stained with DAPI. Scale bars 10 μm unless otherwise specified.

Moreover, the apical localization of both adaptor RNAs showed sensitivity to Puro but not CHX (**Figure 5E,F**), although the change measured in *Bsg25D* signal distribution along the A-B axis in the FE was not significantly different from untreated control (**Figure 5G**). However, *Bsg25D* is expressed at low levels in the FE, hindering a robust quantitative image analysis. To further investigate whether *hook* and *Bsg25D* use the same localization mechanism as *BicD*, we analyzed their spatial relationship with Dhc-GFP/*Dhc* RNA particles. As for *BicD*, both *Bsg25D* (**Figure 5H**) and *hook* (**Figure 5I**) were shown to partially co-localize with, thus decorate Dhc-GFP foci, which also contain *Dhc* RNA. Overall, these results suggest that the RNAs encoding the dynein activating adaptors *BicD*, *hook*, and *Bsg25D*, represent a subgroup of apical RNAs that share the same co-translational, dynein-dependent mechanism that ensures their localization at cortical dynein foci also containing *Dhc* RNA.

## DISCUSSION

Only few examples of localizing RNAs in the FE have been described to date, with little mechanistic insight (Jambor et al., 2015; Li et al., 2008; Horne-Badovinac & Bilder, 2008; Vazquez-Pianzola et al., 2017; Schotman et al., 2008; Serano & Rubin, 2003). To explore the extent of RNA localization in a somatic tissue *in vivo* and gain insight into the mechanisms underlying the phenomenon, we have used laser-capture microdissection of apical and basal subcellular fragments of columnar follicle cells coupled with RNA-seq to identify localizing RNAs in this tissue. This allowed us to investigate in detail the landscape of mechanisms that mediate both apical and basal RNA localization in the FE (**Figure 6A**). In our study, we found that basal RNA localization is mechanistically analogous to posterior RNA localization in the oocyte (represented by *osk*), reflecting MT plus end enrichment (Clark et al., 1997). Khc, *a*Tm1, and the EJC appear to be core components of a general “basal” RNA localization machinery. These results are in line with previous findings on *osk* RNA indicating that Khc/*a*Tm1 bind to the 3’UTR (Gaspar et al., 2017) and the EJC activates kinesin-1 transport through association with the coding sequence (Ghosh et al., 2012).

**Figure 6.**
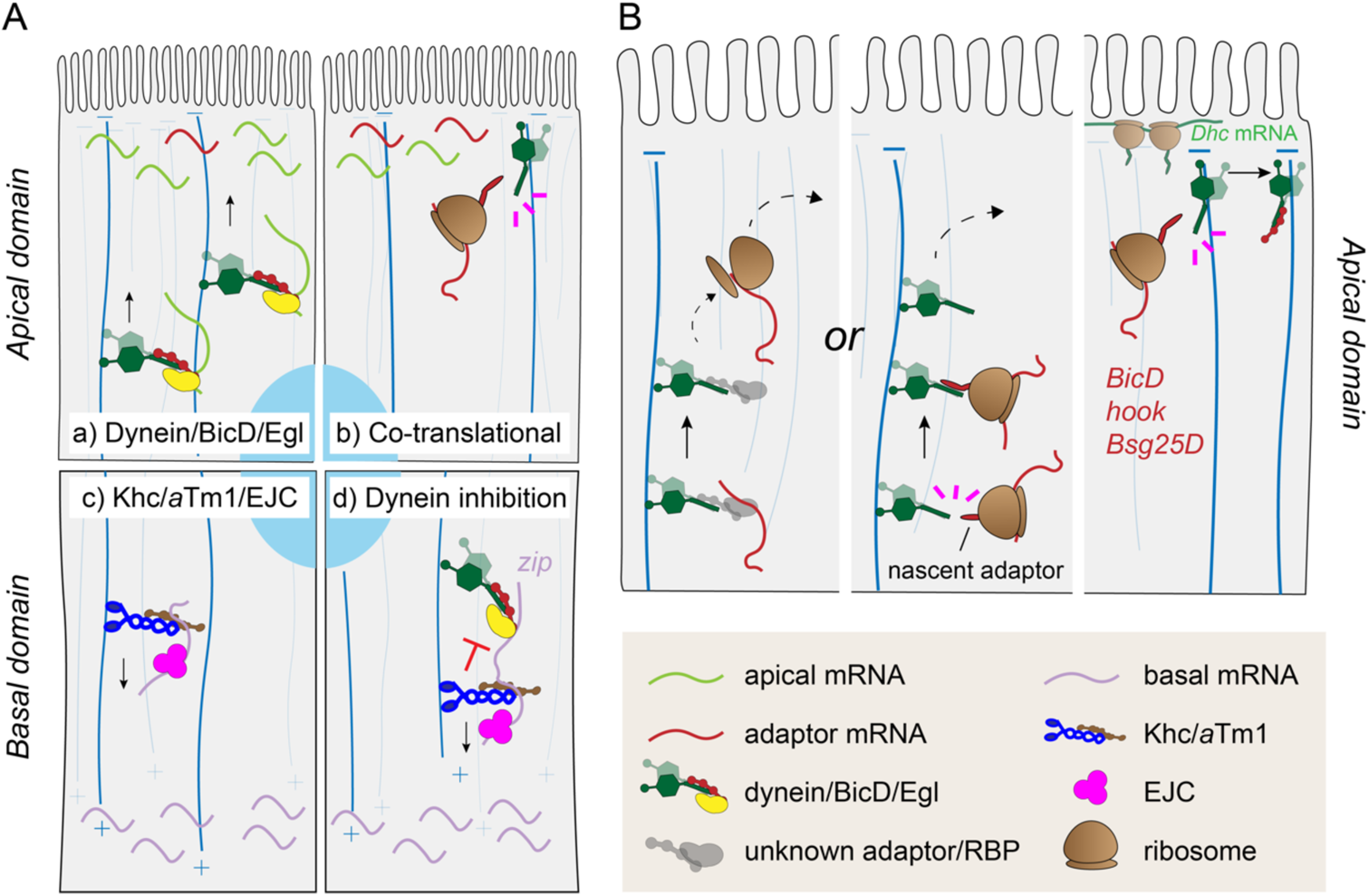
**Models of RNA localization mechanisms in follicle cells.** A) Model of the mechanism underlying apical (a-b) and basal (c-d) RNA localization. Apical RNAs are localized at MT minus ends by two dynein-dependent mechanisms: a) the dynein/BicD/Egl RNA transport machinery localizes most of apical RNAs in the FE; b) a subset of dynein adaptors RNAs (*BicD*, *Bsg25D*, and *hook*) localize co-translationally at cortical dynein foci. c) Basal RNAs are localized by Khc/*a*Tm1/EJC moving towards MT plus ends enriched basally. d) In the transport of basal RNAs, the dynein complex is kept in an inhibited state by kinesin-1 and its regulators. B) Model for the apical localization of adaptor RNAs. RNAs can reach the apical domain by either canonical RNA transport by an unknown RBP complex or by interaction of the nascent adaptor protein with dynein/dynactin transporting the translationally engaged RNA to the apical domain. Once at the apical cortex, the nascent adaptor associates through its N-terminal domain with newly translated cortically-anchored dynein, presumably allowing the relief of both proteins’ autoinhibition.

Interestingly, when either component of the kinesin-1 transport complex was lacking, basal RNAs were mislocalized to the apical domain in a dynein-dependent process. Therefore, dynein-mediated apical localization represents a “default” mechanism that must be overcome by kinesin-1 to drive basal RNA localization. Two possible scenarios could explain dynein-mediated apical mislocalization upon kinesin inhibition. Dynein and kinesin-1 could be engaged in a tug of war, pulling the RNAs in opposing directions, a phenomenon observed in the transport of vesicles and lipid droplets (Hancock, 2014). Alternatively, the dynein complex could be kept in an inhibited state and activated upon disruption of kinesin-1 and its regulators. If the tug-of-war scenario were correct, we would have expected a change in *zip* RNA localization in all RNAi conditions including *egl* RNAi alone, namely a shift to a more basal localization due to the enhanced Khc-dependent motility. However, since we did not see a significant change in *zip* localization when only Egl was knocked down, the tug-of-war hypothesis appears to be less likely than the inhibition hypothesis. In addition, this phenomenon recalls *osk* RNA mislocalization to the oocyte anterior upon disruption of kinesin-1, *a*Tm1 or EJC components (Brendza et al., 2000; Cha et al., 2002; Erdélyi et al., 1995; Hachet & Ephrussi, 2001; Mohr et al., 2001; Newmark & Boswell, 1994; Palacios et al., 2004; Zimyanin et al., 2008) which was hypothesized to occur due to a failure to inactivate dynein-mediated RNA transport (Zimyanin et al., 2008).

Apical RNA localization, on the other hand, can be divided into two mechanistically distinct categories, both based on dynein-mediated transport. The first category includes those RNAs that are transported apically by the dynein/BicD/Egl machinery, a well characterized RNA transport complex that directs RNAs towards MT minus ends in a variety of tissues (Bullock & Ish-Horowicz, 2001). Our data suggest that the majority of apically localizing RNAs may belong to this class, as the localization of most of our randomly chosen apical RNAs was affected in both *Dhc* RNAi and *egl* RNAi conditions. This hypothesis is consistent with previous studies that identified several apical RNAs as BicD/Egl cargoes, in a variety of *Drosophila* tissues (Li et al., 2008; Bhagavatula & Knust, 2021; Karlin-McGinness et al., 1996; Jambor et al., 2014; Vazquez-Pianzola et al., 2017; Van De Bor et al., 2011).

The second category of dynein-dependent apical RNAs does not involve Egalitarian activity for their localization. This includes a subgroup of dynein-activating adaptors, namely *BicD, hook,* and *Bsg25D* (*BICD2, HOOK1-3,* and *NIN/NINL* in mammals). Common features of their apical RNA localization include sensitivity to puromycin and partial co-localization with cortical dynein foci containing also *Dhc* RNA. Puromycin causes the disassembly of the translational machinery and the release of the N-terminal peptides emerging from ribosomes. As the N-terminal portion of these adaptors was shown to bind dynein or dynactin subunits (Hoogenraad et al., 2003; Chowdhury et al., 2015; Urnavicius et al., 2015; Schroeder & Vale, 2016; Redwine et al., 2017; Lee et al., 2020), we propose that the apical localization of *BicD, hook,* and *Bsg25D* depends on the co-translational association between dynein components and nascent adaptors at cortical dynein foci (**Figure 6B**). This process might also be conserved in mammals, since the localization of both *BICD2* and *NIN* RNA was shown to be puromycin-sensitive (Safieddine et al., 2021). Previous studies have shown that dynein/dynactin particles have a low affinity to MTs and predominantly exhibit non-processive movements (Torisawa et al., 2014; Trokter et al., 2012). BICD2, HOOK3 and NIN/NINL were shown to promote the formation of highly processive dynein/dynactin complexes (McKenney et al., 2014; Schalager et al., 2014; Redwine et al., 2017). Therefore, it is possible that co-translational assembly of components of the dynein-adaptor complexes is necessary to overcome dynein auto-inhibition (Torisawa et al., 2014; Zhang et al., 2017). *BicD, hook,* and *Bsg25D* may co-translationally associate with dynein soon after nuclear export of the RNA, promoting its apical transport in a manner similar to what has been proposed for *PCNT* RNA targeting at centrosomes (Sepulveda et al., 2018). Alternatively, since dynein can also function as a MT-tethered static anchor in mid-oogenesis oocytes and follicle cells (Delanoue & Davis, 2005; Delanoue et al., 2007), the interaction between dynein and nascent adaptor proteins could occur after the RNA has reached the cell cortex by dynein-mediated transport. Indeed, puromycin treatment did not completely abolish the apical enrichment of adaptor RNAs, despite causing a marked decrease in their signal close to the apical cortex, where they decorate dynein cortical foci.

*In vitro* studies have shown that full-length BicD/BICD2 adopts an autoinhibitory conformation resulting from CC1/2 folding onto the CTD-containing CC3 (Hoogenraad et al., 2001; Dienstbier et al., 2009; Stuurman et al., 1999). Although the leading hypothesis in the field is that cargo binding to the CTD is responsible for the alleviation of auto-inhibition by freeing up the N-terminal dynein-binding domain (Dienstbier et al., 2009; Hoogenraad et al., 2001, 2003; Matanis et al., 2002), it is possible that *in vivo* both nascent BicD interaction with dynein and cargo binding to the CTD might cooperate in preventing BicD intramolecular inhibition in the cellular environment. Strikingly, whereas the mechanism underlying oocyte localization of *BicD* RNA during mid-oogenesis resembles that observed in follicle cells, the nurse cell-to-oocyte transport of *BicD* RNA appears to be governed by a different, translation-independent mechanism that may not involve interaction with Dhc/*Dhc* RNA particles, consistent with a previous study indicating that *BicD* RNA is translationally inhibited by Me31B in the nurse cells (Nakamura et al., 2001). In contrast to early egg chambers in which the MT network emanates from a posteriorly-positioned microtubule organizing center in the oocyte, mid-stage oocytes and columnar follicle cells are both characterized by non-centrosomal MTs tethered to the cell cortex (Tillery et al., 2018). Therefore, the establishment of ncMTs could be at the basis of the mechanistic switch from translation-independent to co-translational *BicD* RNA localization in these compartments. Strikingly, a recent report has shown that *NIN* RNA (the mammalian ortholog of *Bsg25D*) localizes at ncMTs and its expression is essential for apico-basal MT formation and columnar epithelial shape (Goldspink et al., 2017). Therefore, it is possible that the co-translational transport of adaptor RNAs may be important for correct ncMT nucleation at the apical cortex of the follicular epithelium.

## MATERIALS AND METHODS

### LCM sample preparation

*w1118* virgin females were kept with males for 24 h at 25°C on yeast-supplemented cornmeal food. Ovaries were dissected in PBS, transferred to a cryomold and snap-frozen in cold 2-Methylbutane after removal of excess PBS. Frozen ovaries were immediately covered with OCT cryoembedding compound (Sakura) and snap-frozen again. Before cryostat sectioning, each block was equilibrated at −20°C for 1 h. 10 μm cryosections of OCT-embedded ovaries were carefully placed on a MembraneSlide NF 1.0 PEN (Zeiss), briefly thawed at RT and immediately fixed in 75% RNase-free (RF) ethanol for 30 s. Excess OCT was removed with ddH2O RF, and slides were stained in 100 μl Histogene staining solution (Arcturus) according to the manufacturer’s instructions. Finally, sections were dehydrated in increasing ethanol concentrations (75%, 95%, 100%), and briefly air-dried before LCM.

### LCM and RNA-seq

LCM was performed with a Zeiss PALM MicroBeam and visualized under a 63X objective. Sectioned mid-oogenesis egg chambers were staged according to morphological criteria. Once stage 9-10 egg chambers had been identified, either the apical half (“apical fragment”) or the basal half (“basal fragment”) of 5-10 contiguous columnar follicle cells was microdissected and collected into the cap of an AdhesiveCap tube (Zeiss). 10 fragments of either apical or basal sample type from different egg chambers were pooled for each replicate, with a total microdissected area of ∼3000-4000 μm^2^/replicate. LCM samples were processed according to Chen et al. (2017) to produce high-quality Illumina sequencing libraries. Samples were multiplexed and simultaneously sequenced in a single lane using the NextSeq500 system according to the manufacturer’s instructions.

### RNA-seq analysis

Pre-processing of demultiplexed raw reads was performed on EMBL’s instance of Galaxy platform. Read quality was checked after each processing step with FastQC (Andrews, 2010). Low-quality bases and adapter sequences were trimmed from raw read with Trimmomatic (Bolger et al., 2014). rRNA-filtered reads (SortMeRNA, Kopylova et al., 2012) were mapped against *D. melanogaster* Release 6 (dm6) reference genome with STAR (Dobin et al., 2013). To control for RNA degradation that might have occurred during LCM, the normalized transcript coverage of the uniquely mapping reads was calculated with CollectRNAseqMetrics (part of Picard tools, http://broadinstitute.github.io/picard/). Uniquely mapped reads were counted with featureCounts (Liao et al., 2014) and normalized with DESeq2 (Love et al., 2014). Differential gene expression analysis was performed with DESeq2 by comparing the mean read counts of the Apical (4 replicates, A1-A4) and Basal (4 replicates, B1-B4) samples. Replicates A5 and B5 were excluded from further analysis due to their high degree of dissimilarity with replicates of the same sample type as shown by PCA and Euclidean distance analysis, probably due to a high degree of contamination from neighboring tissues. Statistical significance was set to an FDR-adjusted p value < 0.1 (Benjamini-Hochberg correction for multiple testing). The R package ComplexHeatmap (Gu et al., 2016) was used to generate the heatmap in Figure 2B.

### Identification of contaminant reads

Identification of contaminant RNAs was performed with R Studio. Among the RNAs that were significantly enriched in either the apical (log2FC > 0) or the basal (log2FC < 0) domain, were considered “contaminants” those RNAs displaying high absolute log2FoldChange (|log2FC|), indicating that they were probably originating from neighboring tissues. A threshold of log2FC > 3 and log2FC < −3 was arbitrarily set to identify putative apical and basal contaminants, respectively. The functional annotation of each contaminant candidate was retrieved on FlyBase (release FB2020_6) (Larkin et al., 2021) and their read distribution among apical and basal replicates analyzed through Integrative Genomics Viewer (IGV) (Robinson et al., 2011).

### Fly stocks and genetics

All fly stocks were maintained at 18°C on standard fly food. For crosses, virgin females were mated with *w1118* males at 25°C on cornmeal food supplemented with yeast. Female offspring of the desired genotype were incubated with *w1118* males on a yeast-supplemented medium for 24h at 25°C to stimulate the development of vitellogenic stage egg-chambers before ovary dissection.

The following stocks were obtained from the Bloomington Drosophila Stock Center (BDSC): *w1118* (wild-type; #3605), *DhcRNAi* (#36698), *eglRNAi* (#28969), *KhcRNAi*(#35409), *UAS-NLS-mCherry* (#38425), *osk-Gal4* (#44242), *VK33* (#9750). Other stocks used were: *HsFLP; arm>f+>Gal4; UAS-CD8-mCherry* and *tj-Gal4/CyO* (gifts of Juan Manuel Gomez Elliff), *Tm1^eg1^/TM3Sb,Ser* and *Tm1^eg9^/TM3Sb,Ser* (Erdélyi et al., 1995), *Dhc64C-GFP* (Gáspár et al., 2021), *vasa-Gal4/TM3Sb* (gift of Jean Rene Huynh), *UAS*-*ΔC-Pym-GFP* (Ghosh et al., 2014), *UAS-Egl* (Bullock et al., 2006), *egl^WU50^/SM1* and *egl^PR29^/SM6A* (Mach & Lehmann, 1997). Transgenic flies carrying *UAS-GFP*, *UAS-^0^BicD-GFP*, *UAS-^(+1)^BicD-GFP*, and *UAS-^(−1)^BicD-GFP* were generated in this study by phiC31 integrase-mediated recombination using the VK33 line, which carries an attP site on the third chromosome.

For the generation of *egl^NULL^FC* flies, *egl^WU50^/CyO*; *osk-Gal4/TM3Ser* were crossed with *egl^PR29^/CyO; UAS-Egl/TM3Ser* to generate *egl^WU50^/egl^PR29^; osk-Gal4/UAS-Egl*, expressing Egl only in the germline lineage to rescue the formation of rudimentary ovaries. *tj-Gal4* and *vasa-Gal4* drivers were used to express UAS-containing transgenes in the whole follicular epithelium and in the germline, respectively. To generate flies for FC mutant clone induction, male flies carrying a UAS-containing transgene were crossed with *hsFlp; arm>f+>Gal4; UAS-CD8-mCherry* virgins, and F1 females were subjected to heat-shock as described below.

To generate flies for induction of FC mutant clones in the experiment illustrated in Figure S3, *HsFLP; arm>f+>Gal4/CyO; KhcRNAi/TM6B,Tb* flies were crossed with *+; UAS-NLS-mCherry/CyO; eglRNAi/TM3Ser*. F1 Female flies the desired genotypes [*eglRNAi/TM6B,Tb* for the *egl* RNAi condition; *KhcRNAi/TM3Ser* for the *Khc* RNAi condition; *eglRNAi/KhcRNAi* for the (*egl+Khc)* RNAi condition] were collected and subjected to heat-shock as described below.

### Generation of follicle cell clones

The UAS-Gal4 “flip-out” system was used to generate marked mutant clones in a wild-type background (Struhl & Basler, 1983; Pignoni & Zipurski, 1997). Freshly eclosed females resulting from each cross were collected and mated with *w1118* males for 24 h at 25°C on food supplemented with yeast. Flies were heat-shocked for 1h in a water bath heated at 37°C. According to Gonzales-Reyes & St Johnston (1998), heat-shocked females were kept for 39 h at 25°C with males on yeast before dissection, thus allowing follicle cells that induced the expression of the transgene at stage ∼ 5 to develop into stage 10 follicle cells.

### *Ex vivo* pharmacological treatment

Young *w1118* female flies were incubated with males for 24 h at 25°C on fly food supplemented with yeast. Ovaries were dissected in PBS and immediately incubated in Schneider’s medium (Gibco) supplemented with 15% FBS (Gibco), 0.6X penicillin/streptomycin (Invitrogen), 200 μg/ml insulin (Sigma). For translation inhibitor treatment, either 200 μg/ml puromycin (Gibco) or 200 μg/ml cycloheximide (Sigma) or no compound (control) was added fresh to the medium and ovaries were incubated for 30 min at RT before fixation.

### Generation of BicD-GFP constructs and transgenic fly lines

AttB-pUASp-BicD-GFP-K10 or AttB-pUASp-GFP-K10 plasmids carrying a *w+* cassette, a TLS-deficient version of the K10 3’UTR, and attB sites for phiC31 integrase-mediated recombination into the VK33 line were generated as follows.

To generate plasmid vectors carrying the BicD-GFP gene cassettes (*^0^BicD-GFP, ^(−1)^BicD-GFP*, *^(+1)^BicD-GFP*, *GFP*), BicD and GFP CDS were amplified by PCR and the two fragments were combined into AttB-pUASp-K10 vector by InFusion cloning (Clontech) according to the manufacturer’s instructions. pBS-BicD (BicD-RA, FlyBase ID: FBpp0080555) plasmid (a kind gift from Jean-Baptiste Coutelis) was used as template to generate BicD CDS PCR amplicons. The Fw primer used to amplify BicD CDS was designed in order to include, in addition to a 20 nt-homology with AttB-pUASp-K10 vector, the *Drosophila* Kozak sequence (Cavener, 1987) in frame with a linker sequence where frameshift mutations could be generated, and a region annealing to nt 4-29 of BicD CDS. To generate *^0^BicD-GFP* construct, the 18-bp linker containing the ATG (5’-<u>ATG</u>ATCCTAGGCGCGCGG-3’) was inserted in frame with nt 4-2346 of BicD-RA. To generate *^(+1)^BicD-GFP* construct, a C was inserted at position 4 in the N-terminal 18-bp linker (5’-ATG**<u>C</u>**ATCCTAGGCGCGCGG-3’). To generate *^(−1)^BicD-GFP* construct, a G was deleted at position 10 in the N-terminal 18-bp linker (5’-ATGATCCTA_GCGCGCGG-3’). *^0^BicD-GFP, ^(−1)^BicD-GFP*, and *^(+1)^BicD-GFP* full insert sequences with the respective predicted translated ORF are listed in **File S2**.

To generate UAS-GFP construct, GFP ORF was amplified with a Fw primer containing KpnI restriction site upstream of GFP ATG and with a Rev primer containing NotI restriction site and the stop codon. The amplified fragment was gel purified, digested with KpnI and NotI and ligated into a AttB-pUASp-K10 vector digested with the same enzymes.

Each AttB-containing plasmid was purified and sequenced before injection into VK33 embryos carrying an attP site on the 3^rd^ chromosome. Injected flies were crossed with *If/CyO; Sb/TM3Ser* individuals and transgenic F1 flies were identified by appearance of red eye color.

### Immunostaining

5-10 pairs of ovaries were dissected in PBS and immediately fixed in 2% PFA in PBSTX(0.1%) (PBS + 0.1% Triton-X100) on a Nutator for 20 min at RT, followed by two washes of 15 min each with PBSTX(0.1%) shaking at RT. Ovaries were then blocked in 1X casein/PBSTX(0.1%) (stock: 10X casein blocking buffer, Sigma) for 30 min and incubated with rabbit anti-Egl primary antibody (kind gift from R. Lehmann, Mach & Lehmann, 1997) diluted in blocking buffer o/n at 4°C. Alexa fluor 647 goat anti Rabbit (Jackson Immuno Research) secondary antibody was added in blocking buffer for 2 h at RT. Samples were washed 3x 10 min with 1X casein/PBSTX(0.1%), 1x 10 min with PBSTX(0.1%) + 1:15,000 DAPI and kept o/n in 100 μl of 80% TDE/PBS before mounting on microscope slides.

### Single molecule *in situ* Fluorescence Hybridization (smFISH)

smFISH antisense oligonucleotides (listed in **Table S2**) were designed and labelled with dye-conjugated ddUTPs according to the protocol described by Gáspár et al. (2017) to generate oligonucleotides labelled at their 3’ and with ATTO-633-NHS ester (ATTO-TEC). When dual-color smFISH experiments were performed, each probe set was labelled with either ATTO-633 or ATTO-565. The degree of labelling (DOL, % of labelled oligos) and concentration of the labelled probe sets was measured according to the published algorithm.

Dissected ovaries were immediately fixed in 2% PFA/PBSTX(0.1%) gently shaking for 20 min at RT. In case of *ex vivo* ovary incubation, dissected ovaries were incubated in Schneider’s medium supplemented with the respective pharmacological treatment before proceeding with fixation, as described above. Fixed ovaries were rinsed and washed twice with PBSTX(0.1%) for 10 min before dehydrating them by replacing PBSTX(0.1%) with increasing concentrations of ethanol/PBSTX(0.1%). Fixed and dehydrated ovaries were kept in 100% ethanol at −20°C for up to 10 days until the day of the experiment.

An optimized version of the smFISH protocol described in Hampoelz et al. (2019) was followed with minor modifications. All steps were performed at RT unless specified otherwise. Dehydrated ovaries were first rinsed with PBSTX(0.1%), followed by 2×15 min washes with PBSTX(0.1%), and incubated in Pre-hybridization Buffer (2x SSC, 10% deionized formamide, 0.1% Tween-20) gently shaking for 30 min. The Pre-hybridization Buffer was replaced with 250 μl of Hybridization Buffer (2x SSC, 10% deionized formamide, 0.1% Tween-20, 2 mM vanadyl ribonucleoside complex (New England Biolabs), 100 μg/mL salmon sperm DNA (Invitrogen), 10% dextran sulfate, 20 μg/mL BSA) pre-warmed at 37°C in which smFISH probes were added to a final concentration of 1 nM/probe. Ovaries were kept hybridizing in the dark for 16-17 h on a heat block set at 37°C shaking at 1000 rpm. To remove the excess probes, ovaries where washed 3x 10 min at 37°C with Washing Buffer (2x SSC, 10% deionized formamide, 0.1% Tween-20). 1:15,000 DAPI was added to the second wash. Finally, samples were rinsed 4x in PBST(0.1%) (PBS + 0.1% Tween20) and kept in in 100 μl of 80% TDE/PBS for at least 1 h before mounting on microscope slides.

Z-stacks of images were acquired on a Leica TCS SP8 confocal microscope with 405nm, 488 nm, 552 nm and 640 nm fixed excitation laser lines using a 63X 1.3 NA glycerol immersion objective. A suitable range for spectral detection was carefully chosen for each channel to avoid cross-talk of fluorescence emission. Images were automatically restored by deconvolution with the Lightning module.

### Image analysis and statistical testing

To quantify smFISH fluorescence of localizing RNAs, average Z-projections of deconvolved confocal image stacks were analyzed with Fiji (Schindelin et al., 2012). In mosaic FE, for each wild-type (wt, unmarked) and mutant (mCherry-marked) group of cells within the same Z-stack, a region of interest (ROI) was drawn encompassing the apical and the basal cytoplasm of 5-10 adjacent follicle cells (with the exclusion of nuclei); in addition, a ROI was drawn in an area of the image were no signal was present (background, bg). The mean fluorescence intensity (m.f.i.) was measured for each ROI.

The degree of apicality (DoA) of a given RNA in each cell type (*t*) (wild-type or mutant) and each experimental condition *c*, was measured as follows:

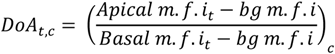

To quantitatively analyze changes in RNA localization in each experimental condition *c*, the DoA measured in mutant (KD) cells was divided by the DoA measured in neighboring wild-type (wt) cells within the same Z-stack:

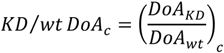

Only in *Tm1^NULL^* condition, due to the impossibility to obtain a mosaic tissue, the DoA of a given RNA in each cell type (*t*) (wild-type or *Tm1^NULL^*), was measured as follows:

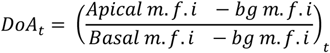

To calculate the change in DoA, the DoA measured in single *Tm1^NULL^* egg chambers was divided by the average DoA measured in *n* wild-type egg chambers (wt):

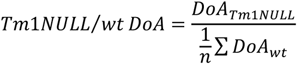

In Figure 2D, Figure 3C, and Figure 5D, KD/wt DoA values of each RNA and experimental condition was measured across at least 3 different Z-projections, and the average value obtained was compared to the null hypothesis *H_0_*: KD/wt DoA=1, corresponding to no change in RNA localization bias following KD treatment [DoA(KD)=DoA(wt)]. One-sample Student’s *t*-test was used to compare means to a reference value of mu = 1 in each experimental condition.

In Figure S3B, independent Student’s *t*-test was used to compare mean DoA(wt) and DoA(RNAi) values in each condition. In Figure S3C, mean KD/wt DoA values across conditions were compared by one-way ANOVA followed by Tukey’s post-hoc tests.

### Other image analysis and statistical procedures

Fluorescence intensity along lines were measured with Fiji on average Z-projections of confocal images and plotted with R Studio. Intensity values from each channel were normalized to 0-1 range. *BicD, GFP, Bsg25D or hook* mean fluorescence intensity along the A-B axis of the epithelium was measured in groups of 5-10 adjacent follicle cells as line plots. At least 3 line plots were generated for each RNA measured in each condition. The value corresponding to 50% of the cumulative area under the curve (a.u.c.) of each plot was considered as the variation of the respective RNA localization along the A-B axis of the epithelium. Welch two sample t-test was used to compare mean values of the 50% of the a.u.c with respect to untreated controls (pharmacological experiments) or in-frame *BicD-GFP* (*Frameshift* vs. *In-frame* variation).

## DATA AVAILABILITY

The authors declare that all data supporting the findings of this study are available within the article and its supplementary information files or from the corresponding author upon reasonable request. RNA-seq data have been deposited in the ArrayExpress database at EMBL-EBI (www.ebi.ac.uk/arrayexpress) under accession number E-MTAB-9127.

## AUTHOR CONTRIBUTIONS

Conceptualization, L.C. and A.E.; Investigation, L.C.; Data Analysis, L.C; Writing – Original Draft, L.C.; Writing – Review & Editing, L.C. and A.E; Supervision, A.E.; Funding Acquisition, A.E.

## ACKNOWLEDGEMENTS

We thank the Bloomington Drosophila Stock Center (BDSC), J. M. Gomez Elliff, A. Debec, I. Gaspar, S. Bullock, J. R. Huynh and R. Lehmann for providing fly lines and reagents. We are grateful to I. Gaspar for helpful discussions and K. Zarnack for training and advice in bioinfomatic analysis. We thank the EMBL GeneCore Facility (D. Pavlinic, V. Benes), Advanced Light Microscopy Facility (S. Terjung), Drosophila Injection Service (A. Reversi), Centre for Bioimage Analysis (C. Tischer), W. Huber and the Centre for Statistical Data Analysis (B. Klaus) and Genome Biology Computational Support (C. Girardot) for their support. We are grateful to Kathi Zarnack, Paolo Ronchi, Luigi Russo, Alessandra Reversi and members of the Ephrussi lab for critically reading the manuscript. This work was supported by EMBL. L.C. was supported by DFG-FOR 2333 grants EP 37/2-1 and EP 37/4-1 from the Deutsche Forschungsgemeinschaft (Germany) to A.E.

