## Supplementary Figures for "Subcellular spatial transcriptomics identifies three mechanistically different classes of localizing RNAs"

2
3

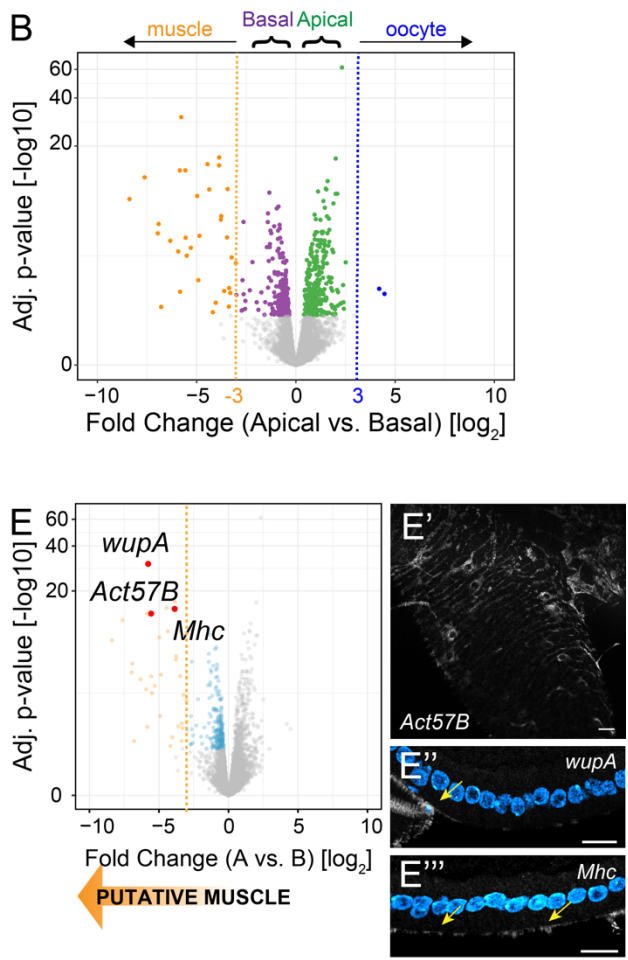

**Figure S1. Identification of LCM contaminant RNAs.** A) Spatial organization of the FE and surrounding tissues. The FE is bounded by the oocyte on its apical side and a thin layer of circular muscle fibers on the basal side. B) Volcano plot depicting apical vs. basal log<sub>2</sub>-transformed fold change (apical vs. basal) values (x axis) by negative log<sub>10</sub>-transformed adjusted p-value (y axis) for all genes analyzed by RNA-seq. Grey points represent genes below the significance threshold, with FDR adj-p value $\geq 0.1$. Significantly enriched genes that exhibit a log<sub>2</sub>FC $< -3$ are highlighted in orange and represent putative muscle contaminants (n=33). Significantly enriched genes that exhibit a log<sub>2</sub>FC $> 3$ are highlighted in blue and represent putative oocyte contaminants (n=2). Green points and purple points represent *bona fide* apical and basal RNAs respectively (FDR adj-p value $< 0.1$, $-3 \leq \log_2\text{FC} \leq 3$). C) Number of reads mapping to significantly enriched genes with log<sub>2</sub>FC $< -3$ (basal contaminants) on log scale. The low mean read count in the apical fragments (green points, avg = 12.5) with respect to the basal fragments (purple points, avg= 336.3) is an indication of contamination of the basal fragments from neighboring tissues and results in the generation of high $|\log_2\text{FC}|$ values. Genes annotated as expressed or having a function in muscle tissue (FlyBase) are highlighted in orange. D) Integrative Genomic Viewer (IGV) representation of reads mapping to the gene model of the basal contaminant *wupA*, with the heatmap showing the number of mapped reads in each apical (n=4, A1-A4) or basal (n=4, B1-B4) sample. E) smFISH validation of *Act57B* (E'), *wupA* (E''), and *Mhc* (E''') RNAs among putative basal contaminants (orange dots, log<sub>2</sub>FC $< -3$ and FDR adj-p value $< 0.1$). Arrows indicate expression of each RNA predominantly in the muscle tissue. The circular fiber organization surrounding each egg chamber can be appreciated by the expression of *Act57B* (top view, E'). Nuclei (cyan) are stained with DAPI. Scale bars 10 $\mu\text{m}$.

Figure S2

A

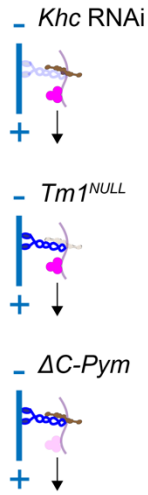

B

*Khc* RNAi mosaic tissue

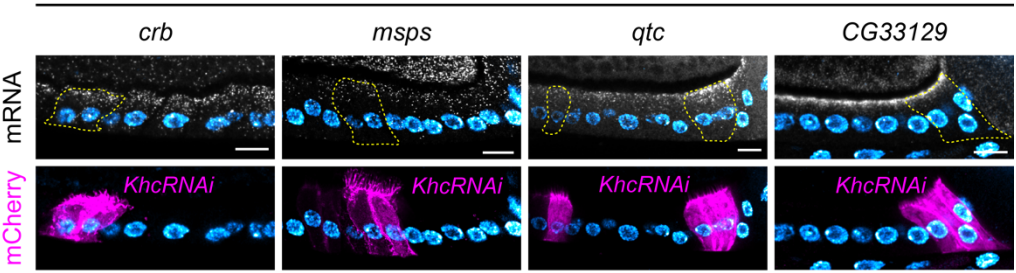

C

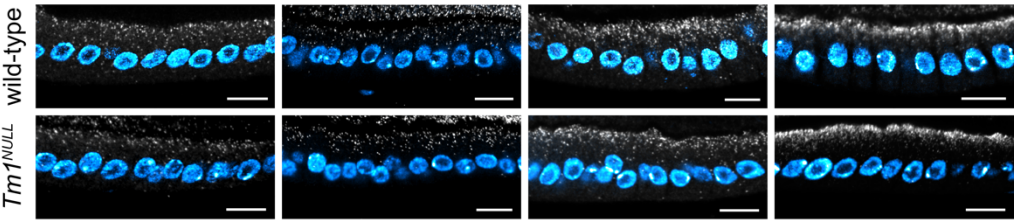

D

ΔC-Pym-GFP mosaic tissue

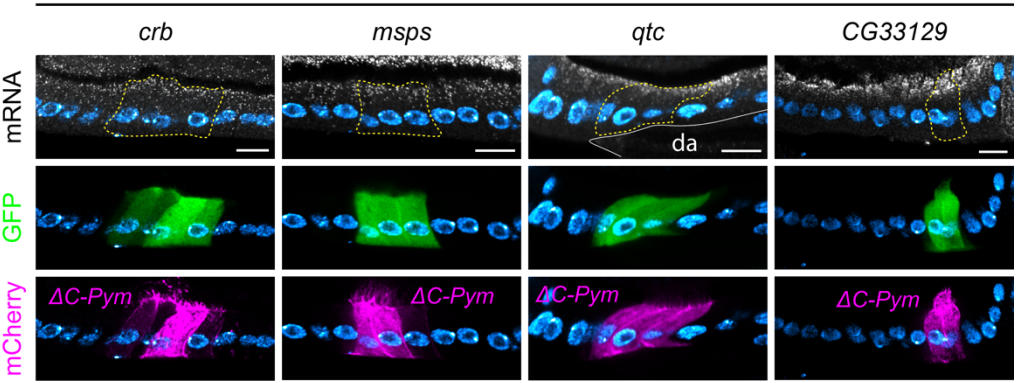

**Figure S2. Effects of kinesin-1 complex disruption in apical RNAs.** A) Schematic representation of RNA transport by Khc/aTm1/EJC. Khc, aTm1, and the EJC were knocked down by *Khc* RNAi, *Tm1<sup>NULL</sup>*, and by expressing $\Delta C$-Pym-GFP, respectively. B) Localization of apical RNAs by smFISH in *Khc* RNAi mosaic tissue. Mutant cells are marked by the expression of CD8-mCherry (lower panels) and highlighted with a dashed line in smFISH images (upper panels). Neighboring wild-type cells are unmarked. C) Localization of apical RNAs by smFISH in wild-type and *Tm1<sup>NULL</sup>* egg chambers. D) Localization of apical RNAs by smFISH in $\Delta C$-Pym-GFP mosaic tissue. Mutant cells are marked by the expression of CD8-mCherry (lower panels) and highlighted with a dashed line in smFISH images (upper panels). Neighboring wild-type cells are unmarked. Nuclei (cyan) are stained with DAPI. Scale bars 10 μm. da = dorsal appendage.

41 **Figure S3**

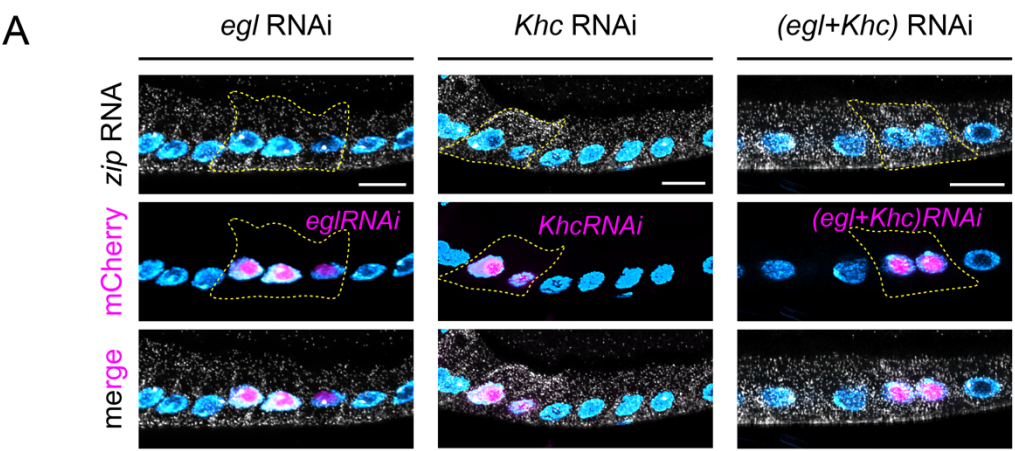

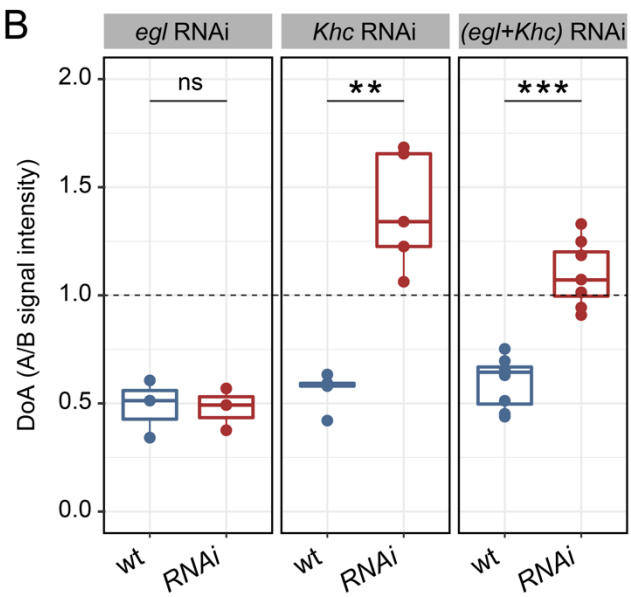

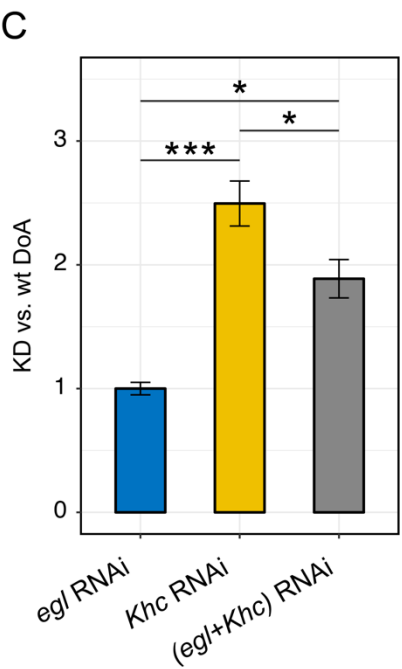

42

43

**Figure S3. Effects of double (*egl+Khc*) RNAi in *zip* RNA localization.** A) *zip* RNA localization in *egl* RNAi, *Khc* RNAi and (*egl+Khc*) RNAi conditions visualized by smFISH. Mutant cells are marked by expression of mCherry in the nucleus (middle panels) and highlighted by a dashed line in smFISH images (upper panels). B) Quantification of *zip* RNA signal (DoA) in wild-type (wt) and RNAi (*RNAi*) cells in *egl* RNAi, *Khc* RNAi and (*egl+Khc*) RNAi conditions. Values of DoA = 1 (dashed horizontal line) correspond to ubiquitous *zip* localization. Student's t test was used to compare means. *egl* RNAi: $p = 0.94$ (ns); *Khc* RNAi: $p = 0.00151$ (\*\*); (*egl+Khc*) RNAi: $p = 0.000004$ (\*\*\*). C) KD/wt change in *zip* DoA shows the variation of *zip* RNA localization in each experimental condition. Values close to 1 indicate no change in *zip* RNA localization upon RNAi. One-way ANOVA ($p = 0.00067$) followed by Tukey post-hoc tests were used to compare means. *egl* RNAi vs. *Khc* RNAi: $p = 0.0005$ (\*\*\*); *egl* RNAi vs. (*egl+Khc*) RNAi: $p = 0.0141$ (\*); *Khc* RNAi vs. (*egl+Khc*) RNAi: $p = 0.0444$ (\*). Nuclei (cyan) are stained with DAPI. Scale bars 10 $\mu\text{m}$.

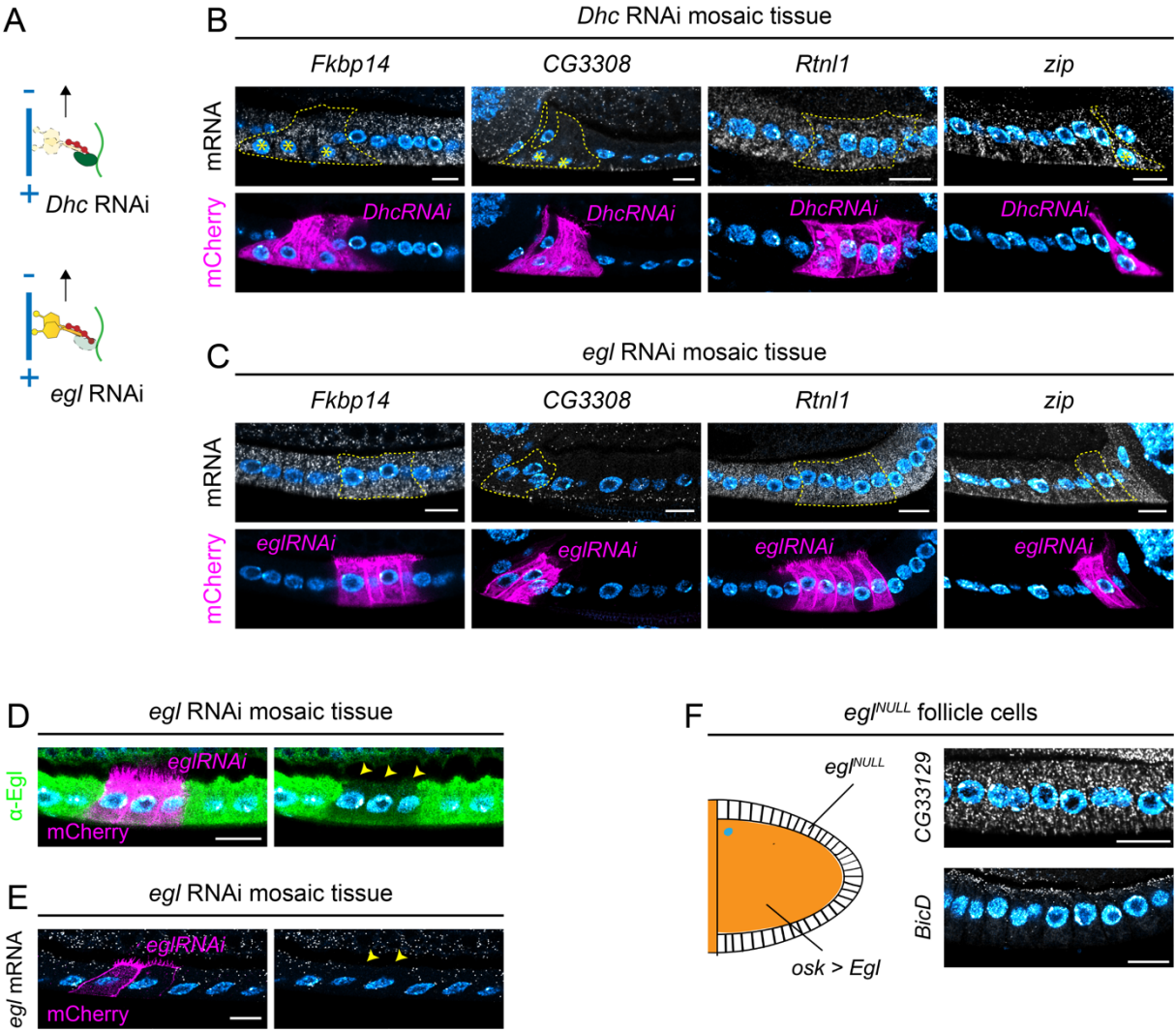

**Figure S4. Effects of dynein/BicD/Egl complex disruption in localizing RNAs.** A) Schematic representation of RNA transport by the dynein/BicD/Egl complex. Dhc and Egl were knocked down by *Dhc* RNAi and *egl* RNAi, respectively. B-C) Localization of basal RNAs by smFISH in *Dhc* RNAi (B) and *egl* RNAi (C) mosaic tissue. Mutant cells are marked by the expression of CD8-mCherry (lower panels) and highlighted with a dashed line in smFISH images (upper panels). Neighboring wild-type cells are unmarked. Asterisks (\*) indicate basal mispositioning of nuclei due to *Dhc* RNAi. D) Expression of Egl protein (green) in mosaic tissue containing wild-type (unmarked) and *egl* RNAi follicle cells (magenta) visualized by immunostaining. Arrowheads indicate the decrease of Egl signal in *egl* RNAi cells. E) Expression of *egl* RNA in a mosaic follicular epithelium containing wild-type (unmarked) and *egl* RNAi cells (magenta) visualized by smFISH using antisense *egl* probes. Arrowheads indicate depletion of *egl* RNA in *egl* RNAi cells. F) *egl*<sup>NULL</sup> egg chambers in which the expression of Egl was rescued only in the germline, resulting in *egl*<sup>NULL</sup> follicle cells (*egl*<sup>NULL</sup>FC, see Materials and Methods for full genotype). In *egl*<sup>NULL</sup>FC, *BicD* RNA was still apically enriched. In contrast, CG33129 RNA was unlocalized, phenocopying *egl* RNAi condition (see Figure 3B). Nuclei (cyan) are stained with DAPI. Scale bars 10 μm.

Figure S5

A

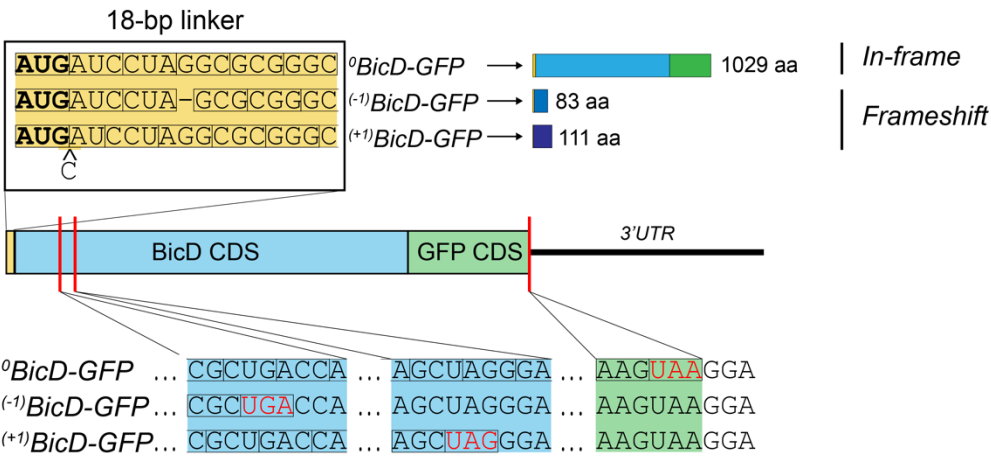

B

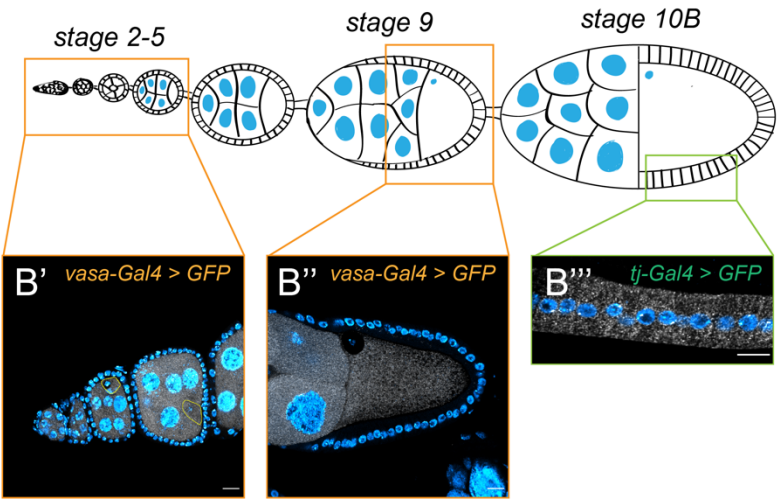

**Figure S5. BicD-GFP and GFP constructs.** A) An 18-bp linker was placed upstream of full-length *BicD* fused in frame with *GFP* to generate frameshift mutations, in an effort to avoid the emergence of RNA localization phenotypes due to the disruption of *BicD* RNA sequence. Each construct differs only in the presence or absence of a single-nucleotide frameshift mutation in the 18-bp N-terminal linker (yellow box). In <sup>0</sup>*BicD-GFP* RNA, the N-terminal linker is translated in frame with *BicD* and *GFP* ORFs and produces a 1029 aa protein that contains a full-length BicD-GFP protein. In <sup>(-1)</sup>*BicD-GFP* RNA, a G was deleted at position 10, causing a -1 frameshift and resulting in a 83 aa peptide. In <sup>(+1)</sup>*BicD-GFP* RNA, a C was added in position 4, causing a +1 frameshift and resulting in a 111 aa peptide. Red lines indicate the predicted termination codons. B) Visualization by smFISH of *GFP* RNA (grayscale), carrying the same 3'UTR as BicD-GFP constructs in the early germline cysts (B'), in mid-stage oocyte (B'') and in the follicular epithelium (B'''). Germline expression was driven by *vasa-Gal4*; expression in the FE was driven by *traffic jam (tj)-Gal4*. Apical is on the top, posterior is on the right. Nuclei (cyan) are stained with DAPI. Scale bars 10 μm.

Figure S6

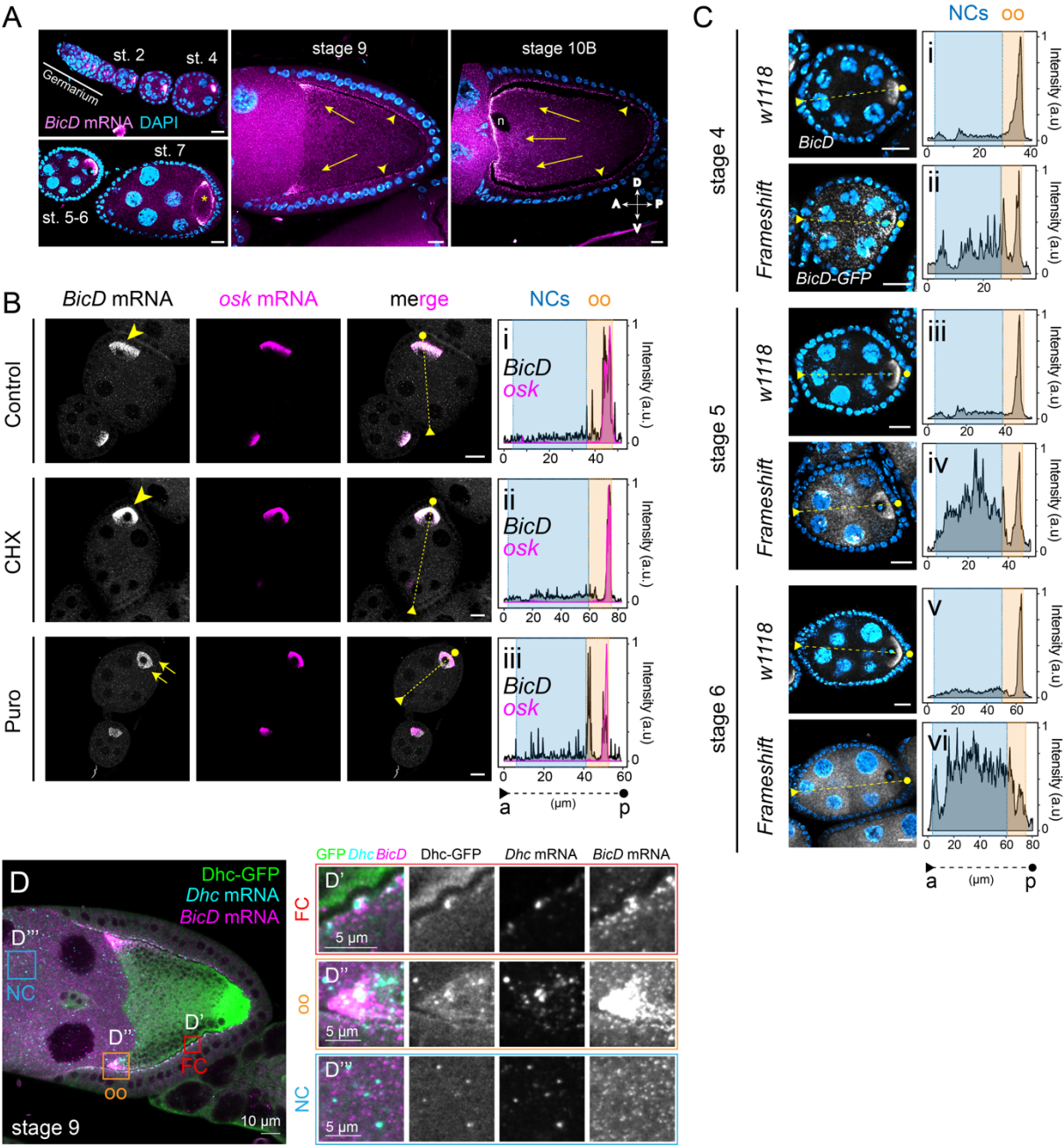

**Figure S6. Translation-dependent and -independent mechanisms of *BicD* RNA localization in the germline.** A) *BicD* RNA localization pattern in throughout oogenesis. *BicD* RNA is enriched in the two pro-oocytes in germarial region 2b, and becomes restricted to the oocyte from stage 1 (germarial region 3) onwards. At stage 5-6 *BicD* accumulates at the posterior of the oocyte. Upon a MT rearrangement event at stage 7 (\*), *BicD* becomes localized at the oocyte corners. At this stage, *BicD* begins to localize apically in the follicle cells surrounding the oocyte. At stage 9, *BicD* localizes at the anterior corners of the oocyte (arrows) and apically in columnar follicle cells (arrowheads). At stage 10, *BicD* is localized at the anterior and lateral cortex of the oocyte (arrows) and appears more tightly apical in the columnar follicle cells (arrowheads). n = oocyte nucleus. A = anterior; P = posterior; D = dorsal; V = ventral. B) Stage 6 egg chambers in control, CHX or Puro conditions. In control and CHX conditions, *BicD* (arrowheads) co-localizes with *osk* at the posterior of the oocyte. Upon Puro treatment, *BicD* RNA becomes ubiquitously distributed in the oocyte (arrows), while *osk* RNA localization is unaffected. Intensity plots represent *BicD* (black) and *osk* (magenta) smFISH signal measured along the A-P axis of each egg chamber on the left. A dashed line indicates the anterior (triangle) to posterior (circle) cross-section along which each fluorescence signal was measured (a.u.). Note the maintenance of *BicD* RNA enrichment in the oocyte in all conditions. C) RNA localization of endogenous *BicD* RNA (w1118) and *Frameshift* *BicD*-GFP RNA in stage 4-6 egg chambers. Intensity plots represent endogenous *BicD* (i,iii,v) or *Frameshift* RNA (ii, iv, vi) smFISH signal measured along the egg chamber A-P axis. Note the ubiquitous distribution of *Frameshift* RNA in the oocyte compared to the posterior localization of endogenous *BicD* RNA. D) Localization of *BicD* RNA, *Dhc* RNA, and endogenously tagged *Dhc*-GFP in stage 9 egg chamber in FC apical cortex (D'), oocyte anterior corners (D''), and nurse cells (D'''). oo = oocyte; FC = follicle cell; NC = nurse cell. a.u. = arbitrary units. a = anterior; p = posterior. In A) and C) nuclei (cyan) are stained with DAPI. Scale bars 10 μm.

**Table S1. List of significantly enriched (FDR < 0.1) RNAs in apical and basal LCM samples.** Apically enriched RNAs (n=306) are characterized by a positive log2FC (log2FC > 0) and are divided into “apical (bona fide)” ($0 < \log_2\text{FC} \leq 3$) or “oocyte contaminant” ($\log_2\text{FC} > 3$). Basally enriched RNAs (n=249) are characterized by a negative log2FC ($\log_2\text{FC} < 0$) and have been divided into “basal (bona fide)” ($-3 \leq \log_2\text{FC} < 0$) or “muscle contaminant” ($\log_2\text{FC} < -3$). Candidate RNAs chosen for validation by smFISH are highlighted in yellow.

**Table S2. List of smFISH oligos used in this study.** Each column contains oligo sequences for all smFISH probe sets used in this study.

**Video S1. Example LCM of basal and apical fragments of a stage 10 follicular epithelium.**

**File S2. Sequence and annotations of BicD-GFP constructs generated in this study.**
