## Supplementary material for "Subcellular spatial transcriptomics identifies three mechanistically different classes of localizing RNAs": File S2

**Supplementary File 2 – Sequences of BicD-GFP constructs**

Legend:

caaaatg Drosophila Kozak sequence

NNNN 18-bp linker

NNNN BicD CDS (excluding ATG and STOP)

NNNN GFP CDS (excluding ATG)

**NNN** predicted STOP codon

_ 1 nt deletion (-1 frameshift)

N 1 nt insertion (+1 frameshift)

**NNNN** predicted translated ORF

>^0^BicD-GFP

caaa**atgatcctaggcgcgggctccagcgccagcaacaacggcccatcggcggaccaatccgtgcaagacctgcaaatggaggtggagcgcctcacgcgcgaactggaccaggtgtcctccgccagcgcccagtccgcccagtacggactgtccctgctggaggagaagtccgccctgcagcagaagtgcgaggaactggagacgctctacgacaacacgcgccacgaactggacatcacacaggaggcgctgaccaagtttcaaacctcacagaaagtgaccaacaagacgggcatcgagcaggaggacgctctgctgaacgaatccgcagctagggagacatcgctcaacctccagatatttgatctggagaacgagcttaagcaactgcgccatgagttggaaagggttcgcaatgagcgcgataggatgctgcaggagaactcggattttgggcgggacaagagcgacagcgaggcggatcgcctacgcctcaagtccgagctgaaggaccttaagttccgggagacgcgtatgcttagcgaatactcggagctggaggaggagaacatatcgctgcaaaagcaggtctccagcctgcgcagctcacaggtggaatttgaaggtgccaaacacgagatccgtcgtctcactgaagaagttgagctgttgaatcaacaggtcgatgagctcgccaatcttaaaaaaattgccgaaaaacaaatggaggaagcgctagagaccttacagggtgaacgtgaggcgaaatatgcattgaagaaggaactggatggccacttgaaccgtgagtctatgtaccacatcagcaacctcgcctacagcatacgcagcaacatggaagacaacgccagcaacaactcggacggtgaggaggaaaatctggctcttaagcgtctggaggctgacctgagcaccgaacttaaatctcctgacggcaccaaatgtgatctcttttcggagattcatctgaacgaactaaagaaactggagaagcagttggagagcatggaaagtgagaagactcatctgacggcgaatttaagggaagctcagacgagtctggacaagtcacaaaacgagctgcagaactttatgtctcgtctggctcttcttgcggcccatgtcgatgctctagtccagctaaagaagcagatcgatgtgaaggagcagggcaaggaaggtggccagaagaaggatgaactggagcagcagctgcgagcgttaatctcgcagtacgccaactggtttacgctctccgccaaggagatcgatggccttaagactgacattgctgaacttcagaagggactcaactatacggacgccaccactacgctgcgcaacgaggtgaccaaccttaagaacaagcttcttgctacggaacaaaagtcactggacctgcagagcgatgttcaaactcttacgcacatctcgcaaaacgctggccaaagtctgggctcagctcgcagtacattggtggccttaagcgacgatctggcgcagctgtatcacctagtttgcacagtcaacggagagacaccgacgcgtgttctgctcgatcataagaccgatgacatgagcttcgaaaacgattctttgactgccatccagtcgcaattcaaatcggatgtctttattgccaagccccagatcgtcgaggatctgcaagggttggcggattccgtggaaattaagaagtacgtggatacagtcagtgatcagatcaagtatctgaagacggctgttgagcataccattgatatgaataaacacaaaatccgctccgagggtggcgacgcactggagaaggttaacacagaggagatggaggaactgcaggagcagatagtcaagttgaagagtttgctgtccgtgaagcgcgagcagattggaactctgcgcaacgtgctcaagtcaaacaagcaaaccgctgaggtggcactgaccaatctcaagtccaagtatgagaacgagaagatcattgtcagcgacaccatgtccaagctacgtaatgagctcaggcttcttaaggaggatgctgccacattctcaagcctgcgtgccatgttcgccgctcgatgcgaggagtatgtgacccaggtggacgatctcaaccgccaattggaggcagcagaggaggagaaaaagactctaaatcagctgttgcgcttggctgtccagcagaagctggcactcactcagcgactcgaggagatggaaatggaccgcgaaatgcgtcacgtccgtcggccgatgccagcccagcgtggcacgagcggcaagtcctccttcagcacgagaccttcgagcaggaatccagcaagcagtaacgccaatccattcgGCGGCCGCgtgagcaagggcgaggagctgttcaccggggtggtgcccatcctggtcgagctggacggcgacgtaaacggccacaagttcagcgtgtccggcgagggcgagggcgatgccacctacggcaagctgaccctgaagttcatctgcaccaccggcaagctgcccgtgccctggcccaccctcgtgaccaccctgacctacggcgtgcagtgcttcagccgctaccccgaccacatgaagcagcacgacttcttcaagtccgccatgcccgaaggctacgtccaggagcgcaccatcttcttcaaggacgacggcaactacaagacccgcgccgaggtgaagttcgagggcgacaccctggtgaaccgcatcgagctgaagggcatcgacttcaaggaggacggcaacatcctggggcacaagctggagtacaactacaacagccacaacgtctatatcatggccgacaagcagaagaacggcatcaaggtgaacttcaagatccgccacaacatcgaggacggcagcgtgcagctcgccgaccactaccagcagaacacccccatcggcgacggccccgtgctgctgcccgacaaccactacctgagcacccagtccgccctgagcaaagaccccaacgagaagcgcgatcacatggtcctgctggagttcgtgaccgccgccgggatcactctcggcatggacgagctgtacaagtaa**

>^(-1)^BicD-GFP

caaa**ATGATCCTA_GCGCGGGCtccagcgccagcaacaacggcccatcggcggaccaatccgtgcaagacctgcaaatggaggtggagcgcctcacgcgcgaactggaccaggtgtcctccgccagcgcccagtccgcccagtacggactgtccctgctggaggagaagtccgccctgcagcagaagtgcgaggaactggagacgctctacgacaacacgcgccacgaactggacatcacacaggaggcgctga**ccaagtttcaaacctcacagaaagtgaccaacaagacgggcatcgagcaggaggacgctctgctgaacgaatccgcagctagggagacatcgctcaacctccagatatttgatctggagaacgagcttaagcaactgcgccatgagttggaaagggttcgcaatgagcgcgataggatgctgcaggagaactcggattttgggcgggacaagagcgacagcgaggcggatcgcctacgcctcaagtccgagctgaaggaccttaagttccgggagacgcgtatgcttagcgaatactcggagctggaggaggagaacatatcgctgcaaaagcaggtctccagcctgcgcagctcacaggtggaatttgaaggtgccaaacacgagatccgtcgtctcactgaagaagttgagctgttgaatcaacaggtcgatgagctcgccaatcttaaaaaaattgccgaaaaacaaatggaggaagcgctagagaccttacagggtgaacgtgaggcgaaatatgcattgaagaaggaactggatggccacttgaaccgtgagtctatgtaccacatcagcaacctcgcctacagcatacgcagcaacatggaagacaacgccagcaacaactcggacggtgaggaggaaaatctggctcttaagcgtctggaggctgacctgagcaccgaacttaaatctcctgacggcaccaaatgtgatctcttttcggagattcatctgaacgaactaaagaaactggagaagcagttggagagcatggaaagtgagaagactcatctgacggcgaatttaagggaagctcagacgagtctggacaagtcacaaaacgagctgcagaactttatgtctcgtctggctcttcttgcggcccatgtcgatgctctagtccagctaaagaagcagatcgatgtgaaggagcagggcaaggaaggtggccagaagaaggatgaactggagcagcagctgcgagcgttaatctcgcagtacgccaactggtttacgctctccgccaaggagatcgatggccttaagactgacattgctgaacttcagaagggactcaactatacggacgccaccactacgctgcgcaacgaggtgaccaaccttaagaacaagcttcttgctacggaacaaaagtcactggacctgcagagcgatgttcaaactcttacgcacatctcgcaaaacgctggccaaagtctgggctcagctcgcagtacattggtggccttaagcgacgatctggcgcagctgtatcacctagtttgcacagtcaacggagagacaccgacgcgtgttctgctcgatcataagaccgatgacatgagcttcgaaaacgattctttgactgccatccagtcgcaattcaaatcggatgtctttattgccaagccccagatcgtcgaggatctgcaagggttggcggattccgtggaaattaagaagtacgtggatacagtcagtgatcagatcaagtatctgaagacggctgttgagcataccattgatatgaataaacacaaaatccgctccgagggtggcgacgcactggagaaggttaacacagaggagatggaggaactgcaggagcagatagtcaagttgaagagtttgctgtccgtgaagcgcgagcagattggaactctgcgcaacgtgctcaagtcaaacaagcaaaccgctgaggtggcactgaccaatctcaagtccaagtatgagaacgagaagatcattgtcagcgacaccatgtccaagctacgtaatgagctcaggcttcttaaggaggatgctgccacattctcaagcctgcgtgccatgttcgccgctcgatgcgaggagtatgtgacccaggtggacgatctcaaccgccaattggaggcagcagaggaggagaaaaagactctaaatcagctgttgcgcttggctgtccagcagaagctggcactcactcagcgactcgaggagatggaaatggaccgcgaaatgcgtcacgtccgtcggccgatgccagcccagcgtggcacgagcggcaagtcctccttcagcacgagaccttcgagcaggaatccagcaagcagtaacgccaatccattcgGCGGCCGCgtgagcaagggcgaggagctgttcaccggggtggtgcccatcctggtcgagctggacggcgacgtaaacggccacaagttcagcgtgtccggcgagggcgagggcgatgccacctacggcaagctgaccctgaagttcatctgcaccaccggcaagctgcccgtgccctggcccaccctcgtgaccaccctgacctacggcgtgcagtgcttcagccgctaccccgaccacatgaagcagcacgacttcttcaagtccgccatgcccgaaggctacgtccaggagcgcaccatcttcttcaaggacgacggcaactacaagacccgcgccgaggtgaagttcgagggcgacaccctggtgaaccgcatcgagctgaagggcatcgacttcaaggaggacggcaacatcctggggcacaagctggagtacaactacaacagccacaacgtctatatcatggccgacaagcagaagaacggcatcaaggtgaacttcaagatccgccacaacatcgaggacggcagcgtgcagctcgccgaccactaccagcagaacacccccatcggcgacggccccgtgctgctgcccgacaaccactacctgagcacccagtccgccctgagcaaagaccccaacgagaagcgcgatcacatggtcctgctggagttcgtgaccgccgccgggatcactctcggcatggacgagctgtacaagtaa

>^(+1)^BicD-GFP

caaa**ATGCATCCTAGGCGCGGGCtccagcgccagcaacaacggcccatcggcggaccaatccgtgcaagacctgcaaatggaggtggagcgcctcacgcgcgaactggaccaggtgtcctccgccagcgcccagtccgcccagtacggactgtccctgctggaggagaagtccgccctgcagcagaagtgcgaggaactggagacgctctacgacaacacgcgccacgaactggacatcacacaggaggcgctgaccaagtttcaaacctcacagaaagtgaccaacaagacgggcatcgagcaggaggacgctctgctgaacgaatccgcagctag**ggagacatcgctcaacctccagatatttgatctggagaacgagcttaagcaactgcgccatgagttggaaagggttcgcaatgagcgcgataggatgctgcaggagaactcggattttgggcgggacaagagcgacagcgaggcggatcgcctacgcctcaagtccgagctgaaggaccttaagttccgggagacgcgtatgcttagcgaatactcggagctggaggaggagaacatatcgctgcaaaagcaggtctccagcctgcgcagctcacaggtggaatttgaaggtgccaaacacgagatccgtcgtctcactgaagaagttgagctgttgaatcaacaggtcgatgagctcgccaatcttaaaaaaattgccgaaaaacaaatggaggaagcgctagagaccttacagggtgaacgtgaggcgaaatatgcattgaagaaggaactggatggccacttgaaccgtgagtctatgtaccacatcagcaacctcgcctacagcatacgcagcaacatggaagacaacgccagcaacaactcggacggtgaggaggaaaatctggctcttaagcgtctggaggctgacctgagcaccgaacttaaatctcctgacggcaccaaatgtgatctcttttcggagattcatctgaacgaactaaagaaactggagaagcagttggagagcatggaaagtgagaagactcatctgacggcgaatttaagggaagctcagacgagtctggacaagtcacaaaacgagctgcagaactttatgtctcgtctggctcttcttgcggcccatgtcgatgctctagtccagctaaagaagcagatcgatgtgaaggagcagggcaaggaaggtggccagaagaaggatgaactggagcagcagctgcgagcgttaatctcgcagtacgccaactggtttacgctctccgccaaggagatcgatggccttaagactgacattgctgaacttcagaagggactcaactatacggacgccaccactacgctgcgcaacgaggtgaccaaccttaagaacaagcttcttgctacggaacaaaagtcactggacctgcagagcgatgttcaaactcttacgcacatctcgcaaaacgctggccaaagtctgggctcagctcgcagtacattggtggccttaagcgacgatctggcgcagctgtatcacctagtttgcacagtcaacggagagacaccgacgcgtgttctgctcgatcataagaccgatgacatgagcttcgaaaacgattctttgactgccatccagtcgcaattcaaatcggatgtctttattgccaagccccagatcgtcgaggatctgcaagggttggcggattccgtggaaattaagaagtacgtggatacagtcagtgatcagatcaagtatctgaagacggctgttgagcataccattgatatgaataaacacaaaatccgctccgagggtggcgacgcactggagaaggttaacacagaggagatggaggaactgcaggagcagatagtcaagttgaagagtttgctgtccgtgaagcgcgagcagattggaactctgcgcaacgtgctcaagtcaaacaagcaaaccgctgaggtggcactgaccaatctcaagtccaagtatgagaacgagaagatcattgtcagcgacaccatgtccaagctacgtaatgagctcaggcttcttaaggaggatgctgccacattctcaagcctgcgtgccatgttcgccgctcgatgcgaggagtatgtgacccaggtggacgatctcaaccgccaattggaggcagcagaggaggagaaaaagactctaaatcagctgttgcgcttggctgtccagcagaagctggcactcactcagcgactcgaggagatggaaatggaccgcgaaatgcgtcacgtccgtcggccgatgccagcccagcgtggcacgagcggcaagtcctccttcagcacgagaccttcgagcaggaatccagcaagcagtaacgccaatccattcgGCGGCCGCgtgagcaagggcgaggagctgttcaccggggtggtgcccatcctggtcgagctggacggcgacgtaaacggccacaagttcagcgtgtccggcgagggcgagggcgatgccacctacggcaagctgaccctgaagttcatctgcaccaccggcaagctgcccgtgccctggcccaccctcgtgaccaccctgacctacggcgtgcagtgcttcagccgctaccccgaccacatgaagcagcacgacttcttcaagtccgccatgcccgaaggctacgtccaggagcgcaccatcttcttcaaggacgacggcaactacaagacccgcgccgaggtgaagttcgagggcgacaccctggtgaaccgcatcgagctgaagggcatcgacttcaaggaggacggcaacatcctggggcacaagctggagtacaactacaacagccacaacgtctatatcatggccgacaagcagaagaacggcatcaaggtgaacttcaagatccgccacaacatcgaggacggcagcgtgcagctcgccgaccactaccagcagaacacccccatcggcgacggccccgtgctgctgcccgacaaccactacctgagcacccagtccgccctgagcaaagaccccaacgagaagcgcgatcacatggtcctgctggagttcgtgaccgccgccgggatcactctcggcatggacgagctgtacaagtaa
